# The DNA replication protein Orc1 from the yeast *Torulaspora delbrueckii* is required for heterochromatin formation but not as a silencer-binding protein

**DOI:** 10.1101/2022.05.06.490984

**Authors:** Haniam Maria, Laura N. Rusche

**Author notes:** Corresponding Author: Address: 109 Cooke Hall, Buffalo, NY, 14260.

## Abstract

To understand the process by which new protein functions emerge, we examined how the yeast heterochromatin protein Sir3 arose through gene duplication from the conserved DNA replication protein Orc1. Orc1 is a subunit of the origin recognition complex (ORC), which marks origins of DNA replication. In *Saccharomyces cerevisiae*, Orc1 also promotes heterochromatin assembly by recruiting the structural proteins Sir1-4 to silencer DNA. In contrast, the paralog of Orc1, Sir3, is a nucleosome-binding protein that spreads across heterochromatic loci in conjunction with other Sir proteins. We previously found that a non-duplicated Orc1 from the yeast *Kluyveromyces lactis* behaved like ScSir3 but did not have a silencer-binding function like ScOrc1. Moreover, *K. lactis* lacks Sir1, the protein that interacts directly with ScOrc1. Here, we searched for the presumed intermediate state in which non-duplicated Orc1 possesses both the silencer-binding and spreading functions. In the non-duplicated species *Torulaspora delbrueckii*, which has an ortholog of Sir1 (TdKos3), we found that TdOrc1 spreads across heterochromatic loci independently of ORC, as ScSir3 and KlOrc1 do. This spreading is dependent on the nucleosome binding BAH domain of Orc1 and on Sir2 and Kos3. However, TdOrc1 does not have a silencer-binding function: *T. delbrueckii* silencers do not require ORC binding sites to function, and Orc1 and Kos3 do not appear to interact. Instead, Orc1 and Kos3 both spread across heterochromatic loci with other Sir proteins. Thus, Orc1 and Sir1/Kos3 originally had different roles in heterochromatin formation than they do now in *S. cerevisiae*.

## INTRODUCTION

Gene duplication enables functionally distinct proteins to emerge without loss of ancestral functions. For multifunctional proteins, a common outcome after duplication is subfunctionalization, in which the ancestral functions are partitioned between the two duplicated copies (Force *et al*. 1999; Conant and Wolfe 2008; Froyd and Rusche 2011). In some cases, subfunctionalization is followed by further change in one or both paralogs, which allows for optimization of activities that were constrained within the multifunctional ancestor (Hittinger and Carroll 2007). Although various models have been proposed to account for the evolutionary trajectories after duplication, less attention has been given to the molecular events that give rise to multifunctional proteins in the first place. Here, we searched for an example of a multifunctional protein that is predicted to be an intermediate in the evolution of the heterochromatin protein Sir3 from the DNA replication protein Orc1.

In the budding yeast lineage, *ORC1* gave rise to *SIR3* after a whole genome duplication approximately 100 million years ago (Wolfe and Shields 1997; Shen *et al*. 2018). Both Orc1 and Sir3 contribute to heterochromatin formation in the well-studied yeast *Saccharomyces cerevisiae*. This heterochromatin is composed of a complex of Sir proteins (Silent Information Regulator) that bind nucleosomes and spread across several kilobases of DNA, resulting in the repression of transcription. Sir heterochromatin forms at the silent mating-type loci, which encode extra copies of mating-type genes that must be repressed to maintain proper cell-type identity. Sir heterochromatin also forms near telomeres, where it is thought to stabilize the ends of chromosomes.

Orc1 is the largest subunit of the origin recognition complex (ORC), which marks origins of DNA replication (Bell and Stillman 1992). In *S. cerevisiae*, ScOrc1 also helps establish heterochromatin at the silent mating-type loci. As part of ORC, Orc1 binds to DNA sequences called silencers and then recruits the Sir proteins. Specifically, Orc1 interacts with Sir1 (Triolo and Sternglanz 1996; Gardner *et al*. 1999), which in turn binds other Sir proteins. In contrast, the paralog of Orc1, Sir3, is a nucleosome-binding protein that spreads across the silent mating-type loci in conjunction with Sir2 and Sir4 (Hoppe *et al*. 2002; Luo *et al*. 2002; Rusche *et al*. 2002; Onishi *et al*. 2007). Orc1 and Sir3 have similar domain structures, with an N-terminal BAH (Bromo Adjacent Homology) domain and a C-terminal AAA+ domain, separated by a disordered linker (Li *et al*. 2018). The BAH domain of ScOrc1 specifically binds ScSir1 (Hou *et al*. 2005; Hsu *et al*. 2005) and is therefore critical for Orc1’s role as a silencer-binding protein. For ScSir3, the BAH domain binds nucleosomes and is critical for the Sir proteins to spread (Buchberger *et al*. 2008; Armache *et al*. 2011). The ScOrc1 BAH domain also binds nucleosomes (De Ioannes *et al*. 2019), but this property is thought to be more important for origin binding than heterochromatin formation.

We previously showed that *ORC1* and *SIR3* subfunctionalized after duplication (Hickman and Rusche 2010). We found that in a non-duplicated yeast species, *Kluyveromyces lactis*, KlOrc1 behaves like ScSir3, facilitating the spreading of Sir proteins through nucleosome binding. Thus, Orc1 already had Sir3-like behavior prior to duplication. Additionally, we found that after duplication the function of the Sir3 AAA+ base subdomain changed, presumably leading to optimization of its ability to promote heterochromatin assembly (Hanner and Rusche 2017). However, one puzzle that remains is when Orc1 acquired a role as a silencer-binding protein. We found that KlOrc1 does not act as a silencer binding protein (Hickman and Rusche 2010). Moreover, *K. lactis* lacks Sir1, which acts as a bridge between ScOrc1 and the Sir proteins and would presumably be required for Orc1 to act as a silencer-binding protein. Thus, the silencer-binding role of Orc1 likely arose after or in conjunction with the emergence of Sir1.

The first recognizable *SIR1* gene appeared more recently than the ancestors of the other *SIR* genes. The appearance of *SIR1* slightly pre-dates the whole-genome duplication, as homologs are found in the clade of non-duplicated yeasts that shares most sequence similarity to duplicated species (Figure 1A). Over evolutionary time, the *SIR1* gene has undergone expansions and contractions in copy number, likely facilitated by its subtelomeric location in many species (Gallagher *et al*. 2009). For example, among duplicated yeast species, *S. cerevisiae* has a single *SIR1* gene, *Saccharomyces bayanus* has four functional *SIR1* homologs (called *KOS* or Kin Of Sir1), and *Candida glabrata* has no *SIR1* homologs (Gallagher *et al*. 2009). In addition to variation in copy number, the *SIR1* gene has undergone an internal tandem duplication. The C-terminal portion of the protein consists of an ORC Interacting Region (OIR) that binds to Orc1 (Triolo and Sternglanz 1996; Gardner *et al*. 1999). The N-terminal portion has a similar sequence, called the OIR’, whose function remains unclear (Connelly *et al*. 2006; Hou *et al*. 2009). Only a single OIR occurs in the presumed ancestral version of the protein, called Kos3. All non-duplicated species that have a Sir1 ortholog have Kos3 but no other Sir1-like proteins (Gallagher *et al*. 2009; Ellahi and Rine 2016).

**Figure 1:**
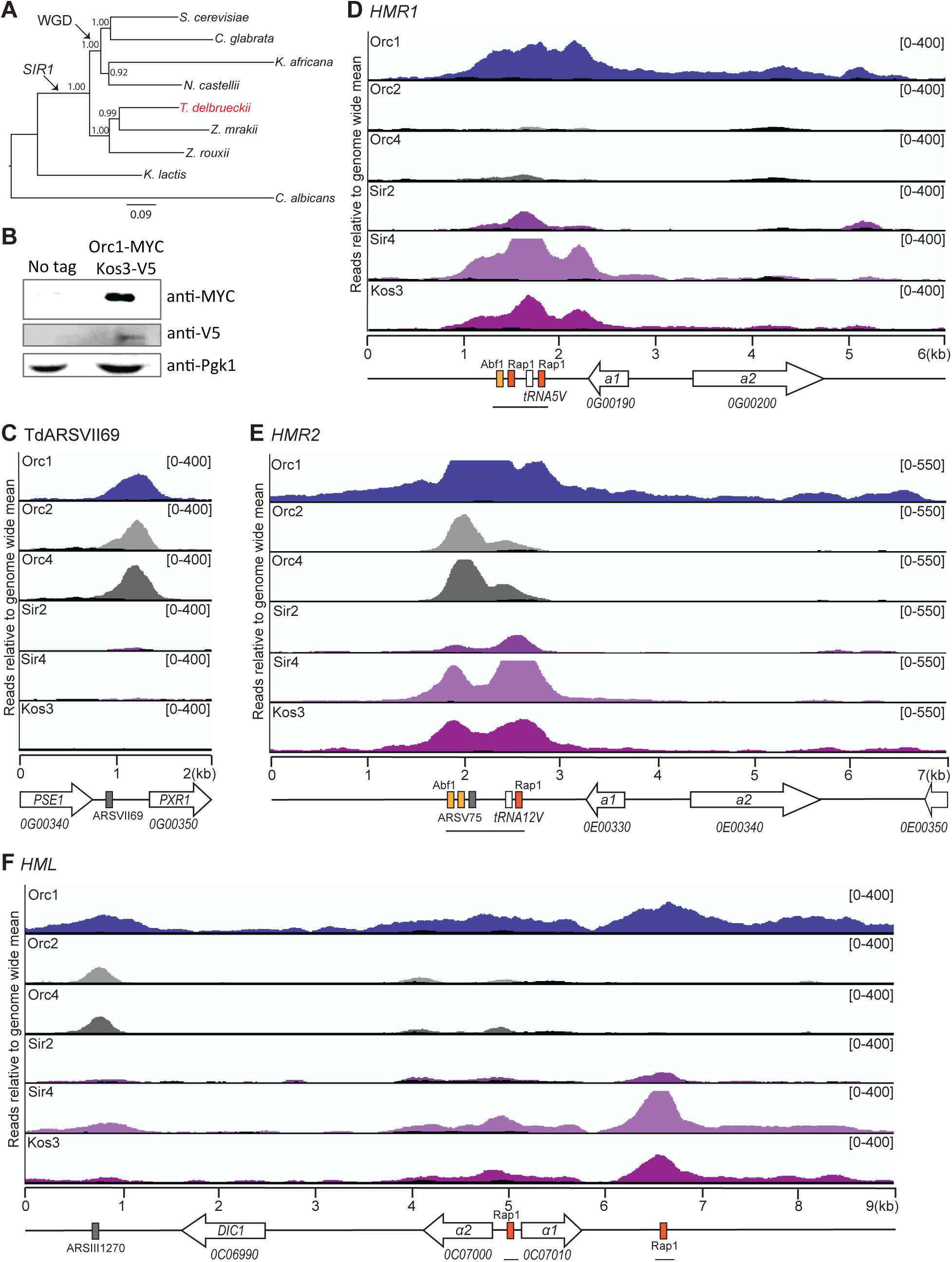
TdOrc1 associated with the mating-type loci and origins of replication. **(A)** Evolutionary relationship of yeast species discussed. WGD denotes whole genome duplication. *SIR1* indicates the appearance of the *SIR1*/*KOS* gene family. The tree was built using 10 genes found in single copy in all species (Aguileta *et al*. 2008). The protein sequences were concatenated and aligned in MAFFT using E-INS-I model, and a maximum likelihood tree was constructed with PhyML using the LG+F+I+G4 model. Node support was assessed with approximate likelihood ratio tests (aLRT). **(B)** Immunoblot of TdOrc1-Myc and TdKos3-V5 (LRY3211). The parent strain (JRY10156) was included as a no tag control. Endogenous 3-phosphoglycerate kinase (Pgk1) was detected as a loading control. **(C-F)** The distribution of TdOrc1-Myc (LRY3211), TdKos3-V5 (LRY3211), TdSir2-V5 and TdSir4-V5 (Ellahi and Rine 2016), and TdOrc2-V5 and TdOrc4-V5 (Maria *et al*. 2021) at **(C)** a replication origin ARS-VII69, **(D)** *HMR1*, **(E)** *HMR2*, and **(F)** *HML* were determined using chromatin immunoprecipitation followed by sequencing (ChIP-Seq). Each track represents the read depth of ChIP-Seq sequences from a single sample. Replicate samples gave similar results. To assess potential variation in sequencing across the genome, read depth for either input chromatin (Orc1, Kos3, Sir2, Sir4) or mock IP (Orc2, Orc4) is overlaid in black. The genomic features are shown below the x-axis. Silencers are marked with solid black lines. Predicted binding sites for potential silencer-binding proteins are represented as boxes for Rap1 (orange), Abf1 (yellow) and ORC (grey).

In addition to having appeared more recently than other Sir proteins, Sir1 differs functionally from the other Sir proteins, at least in *S. cerevisiae*. ScSir1 promotes the establishment of heterochromatin by recruiting the other Sir proteins to the silencer. However, it does not spread across the silent mating-type loci with the other Sir proteins and thus is not required to maintain silencing (Rusche *et al*. 2002; Ellahi and Rine 2016). Consequently, *sir1Δ S. cerevisiae* cells can exist in two epigenetically stable states, in which the mating-type loci are either silenced or expressed (Pillus and Rine 1989). Unlike ScSir1, TdKos3 is not restricted to the silencers and is required for silencing (Ellahi and Rine 2016). Therefore, despite being homologous proteins, ScSir1 and TdKos3 make different contributions to heterochromatin. This different behavior could be related to the presence of a single Orc1 that potentially behaves both as a silencer-binding protein and a spreading protein and therefore recruits Kos3 both to silencers and to sites of spreading.

To search for an ancestral Orc1 that has both silencer-binding and spreading functions, we investigated the properties of Orc1 in *Torulaspora delbrueckii*. This species belongs to the *Zygosaccharomyces-Torulaspora* (ZT) clade of budding yeast (Figure 1A), which has the most sequence similarity to duplicated species and encodes the Sir1 ortholog Kos3. Thus, *T. delbrueckii* could represent an intermediate stage in the evolution of Sir silencing, in which Orc1 already had two distinct roles in heterochromatin formation. However, we found that despite *T. delbrueckii* having a Sir1-like protein, TdOrc1 does not act as a silencer binding protein. In contrast, it does serve as a nucleosome-binding protein that spreads across the heterochromatic loci independently of the ORC complex. Given the absence of ORC binding sites at silencers, we also investigated which proteins do bind these sequences. We found that, Rap1 and Abf1, which bind to *S. cerevisiae* silencers, are also present at *T. delbrueckii* silencers. In contrast, proteins that bind to *K. lactis* silencers were not associated with *T. delbrueckii* silencers. Taken together, our data indicate that a silencer binding function for Orc1 does not go hand-in-hand with a *SIR1* homolog.

## MATERIALS AND METHODS

### Yeast strain construction

Yeast strains used in this study were derived from JRY10156 (Ellahi and Rine 2016), a *Torulaspora delbrueckii MAT*α *ura3Δ0 trp3-1(G466A)* strain descended from NRRL Y-866. Yeast strains are listed in the Reagent Table, plasmids used for strain construction are described in Table S1 and the Reagent Table, and oligos used for strain construction are listed in Table S2. The details of strain construction are provided in the supplemental materials.

### Yeast growth and transformation

Yeast were grown at 30°C in YPD (1% yeast extract, 2% peptone, 2% glucose) or YM (0.67% yeast nitrogen base without amino acids, 2% glucose). *T. delbrueckii* cells were transformed by electroporation (Hickman and Rusche 2009), and *S. cerevisiae* cells were transformed using polyethylene glycol and lithium acetate (Schiestl and Gietz 1989). Transformation details are provided in the supplemental materials.

### Silencing assay using *URA3* reporter

To assess transcriptional silencing, a previously described (Ellahi and Rine 2016) reporter plasmid was used, in which the entire *HMR2* locus was cloned onto a plasmid with a *T. delbrueckii* centromere, an *S. cerevisiae* origin, and a hygromycin resistance gene. To evaluate silencing, the **a**1 open reading frame was replaced with *URA3*. We replaced the hygromycin resistance gene with a kanamycin resistance gene using NEBuilder HiFi DNA assembly master mix to create pLR1377. This plasmid was then modified (Table S3) by replacing the *HMR2* silencer with the corresponding region from *HMR1* (pLR1384), deleting the ORC binding motif (pLR1382), deleting the ORC binding motif plus the Abf1 binding sites (pLR1399), or deleting the *S. cerevisiae* derived plasmid origin (pLR1400 and pLR1401).

To assess silencing of the *URA3* reporter, yeast were transformed with the reporter plasmids described above. Transformed cells were grown overnight at 30°C in 3 mL YPD with 0.3 mg/mL geneticin. Next day, 1 OD_600_ equivalent of cells was harvested and resuspended in 100 µL YM. Five 10X serial dilutions were prepared, and 3 μL of each dilution was plated on CSM (YM supplemented with amino acids and nutrients Sunrise Science 1001), CSM-Ura (YM supplemented with amino acids and nutrients but lacking uracil; Sunrise Science 1004) and CSM+5FOA (YM supplemented with 0.1% 5-fluoroorotic acid, 0.005% uracil, 0.01% tryptophan, and pH adjusted to 3.35). All three media were supplemented with 0.3 mg/mL geneticin. Plates were imaged after 3 days of incubation at 30°C.

### Chromatin immunoprecipitation

Chromatin immunoprecipitation was performed as previously described (Rusche and Rine 2001). Cells were grown in YPD at 30°C, harvested at an OD_600_ of ∼1, and cross-linked for 1 hour in 1% formaldehyde. For immunoprecipitation, 4 µl of anti Myc antibody (Millipore 06-549) or anti V5 antibody (Millipore AB3792) were used. The immunoprecipitation was conducted with 10 μL of Protein A agarose beads. For qPCR analysis, immunoprecipitation was conducted in the presence of 0.25 mg/mL BSA and 0.1 mg/mL salmon sperm DNA. For ChIP-Seq analysis immunoprecipitation was conducted without BSA or salmon sperm DNA. Following immunoprecipitation, samples were reverse crosslinked overnight at 65°C and digested with proteinase K at 45°C.

For ChIP-Seq, library preparation (Takara Bio SMARTer ThruPlex DNA seq kit) and sample barcoding was done at the Next-Generation Sequencing facility at University at Buffalo. The samples were then sequenced on an Illumina 1.9 sequencer using 76 bp single-end sequencing.

For ChIP-qPCR, samples were quantified relative to a standard curve prepared from input DNA. Then, the ratio of the experimental locus to the control locus (*ATS1*, TDEL0H03720) was calculated and presented as relative enrichment. Oligonucleotides used for qPCR are listed in Reagent Table.

### Pipeline for processing ChIP-Seq data

Raw single-end reads were aligned to the *T. delbrueckii* genome (CBS1146 (GCA_000243375.1)) (Gordon *et al*. 2011) using Bowtie2/2.3.4.1 (Langmead *et al*. 2009). Samtools/1.9 was used to convert .sam output files from Bowtie2/2.3.4.1 to .bam, -sorted.bam and index files (Li *et al*. 2009). To identify statistically significant Orc1 and Kos3 genome wide association over background, **M**odel-based **A**nalysis of **C**hIP-**S**eq, MACS2/2.2.7.1 was used to call peaks using one million randomly selected high quality reads with genomic input as control for all samples with the parameters -B, --nomodel, --keep-dup 3, - -extsize 200 (Zhang *et al*. 2008). For visualization bedGraph files generated from MACS2 were processed using bedClip against chromosome sizes extracted from ASM24337v1 (Gordon *et al*. 2011) and converted to .bigwig files using bedGraphToBigWig (Kent *et al*. 2010). ChIP-Seq bigwig files were visualized on IGV (Robinson *et al*. 2011).

### Bioinformatic analysis of Kos3 sequences

Sequences of Kos3 proteins were obtained from NCBI using tblastn to search whole-genome shotgun sequences of Saccharomycetaceae species. For each genus, Kos3 sequences from representative species were used as queries. OIR domains (light grey boxes) were identified by submitting each protein as a query to NCBI blastp. The program returned the boundaries of the “Sir1 superfamily” domain that corresponds to the OIR. Other conserved regions (colored boxes) were identified using the MEME suite (Bailey *et al*. 2009) to search for sequence motifs present zero or one time (ZOOPS model) in each member of the protein set. Search parameters were to search for six motifs per sequence, with a minimum of six and a maximum of 50 amino acids per motif. A zero order Markov model of the background was used.

### Identification of binding sites in silencers

The recognition sequences for Rap1, Abf1, and ORC were predicted at each of the mating-type loci (Table 1) and are indicated by colored boxes in schematics of chromosomal features. For ORC, genome-wide binding sites and motifs were previously reported (Maria *et al*. 2021). For Rap1, we identified motifs associated with ChIP-Seq peaks (unpublished) and then located matches to these motifs within mating-type loci. For Abf1, we identified matches to the known binding site for *S. cerevisiae* Abf1, 5’-TnnCGTnnnnnnTGAT (Beinoraviciute-Kellner *et al*. 2005).

**Table 1:**
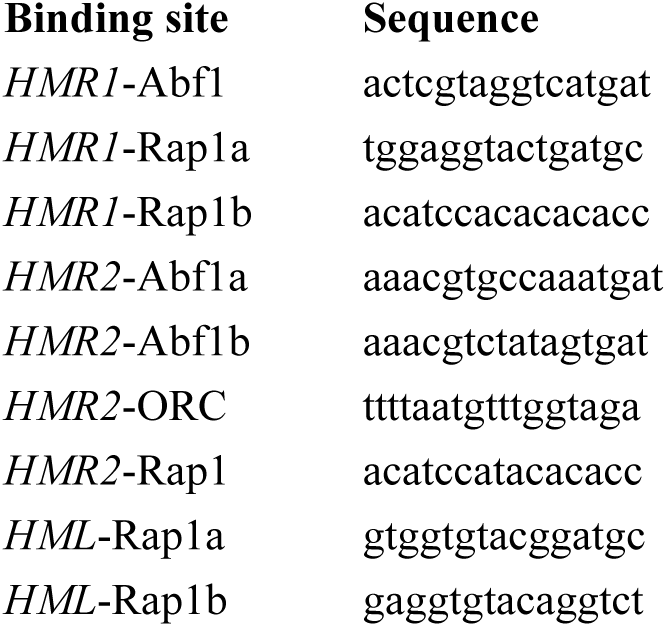
Presumed binding sites at mating-type silencers.

### Construction of species tree

To construct the species tree, we selected 10 protein-coding genes (Table S4) found in single copy across the nine species (Aguileta *et al*. 2008). The protein sequences were obtained from the yeast gene order browser (Byrne and Wolfe 2005) and the candida gene order browser (Maguire *et al*. 2013) and then concatenated, resulting in a combined alignment of 10092 positions. The concatenated sequences were aligned in MAFFT v7.490 using E-INS-i with default settings (Katoh and Standley 2013). MAFFT alignment was input into IQTree (Trifinopoulos *et al*. 2016) to determine the best amino acid substitution model: LG+F+I+G4 (Hoang *et al*. 2018). Finally, a maximum likelihood phylogenetic tree was constructed using PhyML v3.1 using the LG+F+I+G4 model as implemented in Seaview v5.0.5 (Gouy *et al*. 2010). Node support was assessed with approximate likelihood ratio tests (aLRT). Subtree pruning and regrafting (SPR) was used as a search algorithm with five random starts.

## DATA AVAILABILITY

Strains and plasmids are available upon request. The raw Illumina reads for Orc1 and Kos3 ChIP-sequencing and related data that underly this article were uploaded to the NCBI GEO database (Accession number: GSE182111). Supplemental files available at FigShare contain additional methods and a reagent table.

## RESULTS

### TdOrc1 associated with the silent mating-type loci as well as replication origins

To investigate whether Orc1 contributes to the formation of heterochromatin, we determined its distribution across the *T. delbrueckii* genome using chromatin immunoprecipitation followed by high throughput sequencing (ChIP-Seq). For this purpose, we generated a tagged strain with *ORC1*-Myc and *KOS3*-V5 integrated at their genomic loci. Both proteins were detectable by immunoblotting (Figure 1B). Next, we immunoprecipitated and sequenced chromatin fragments associated with Orc1 and Kos3. We also sequenced input chromatin as a control for regions over-represented in the starting material. For comparison, we examined previously published ChIP-Seq data for Sir2, Sir4, Orc2 and Orc4 (Ellahi and Rine 2016; Maria *et al*. 2021). We found that TdOrc1 was associated with all three silent mating-type loci: *HMR1* (chromosome VII), *HMR2* (chromosome V), and *HML* (chromosome III). At these loci, Orc1 was broadly distributed, spanning ∼1.5 kb and overlapping the predicted silencers (Figure 1D-F). Note that *T. delbrueckii* has two *HMR* loci that are nearly identical across the **a**1 and **a**2 genes but different in the silencer regions. The pattern of enrichment for Orc1 was similar to that of the Sir proteins (Sir2, Sir4, and Kos3) at all three loci. In contrast, the Orc2 and Orc4 subunits were absent from *HMR1* and *HML* and only associated with *HMR2* at the previously described ORC binding sequence, ARS-V75 (Maria *et al*. 2021). For comparison, at a representative ORC binding site not at a mating-type locus, TdARS-VII69 (Maria *et al*. 2021), all three ORC subunits were enriched but the Sir proteins were not (Figure 1C). These findings indicate that TdOrc1 has an ORC-independent role at the silent mating-type loci, where it colocalizes with other Sir proteins.

### TdSir2 and TdKos3 facilitated the recruitment and spreading of TdOrc1 at *HMR1* and *HMR2*

In *S. cerevisiae*, ScSir3 requires ScSir2 and ScSir4 to spread across the mating-type loci (Hoppe *et al*. 2002; Rusche *et al*. 2002). To determine if the association of TdOrc1 with the silent mating-type loci also depends on the Sir proteins, we first investigated whether Orc1 associated with *HMR* in the absence of Sir2 by conducting ChIP-Seq on Orc1-Myc in a *sir2Δ* strain. We found TdOrc1 association at *HMR1* was substantially reduced in the absence of Sir2 (Figure 2A, right panel). In contrast, Orc1 association at the nearby origin, ARS-VII22, was not affected (Figure 2A, left panel). Similarly, at *HMR2* Orc1 association was reduced in the absence of Sir2, although it remained associated with the ORC binding site ARS-V75 (Figure 2C, track 3). Thus, Sir2 was required for recruitment of Orc1 to *HMR1* and spreading of Orc1 beyond the ORC binding site at *HMR2*.

**Figure 2:**
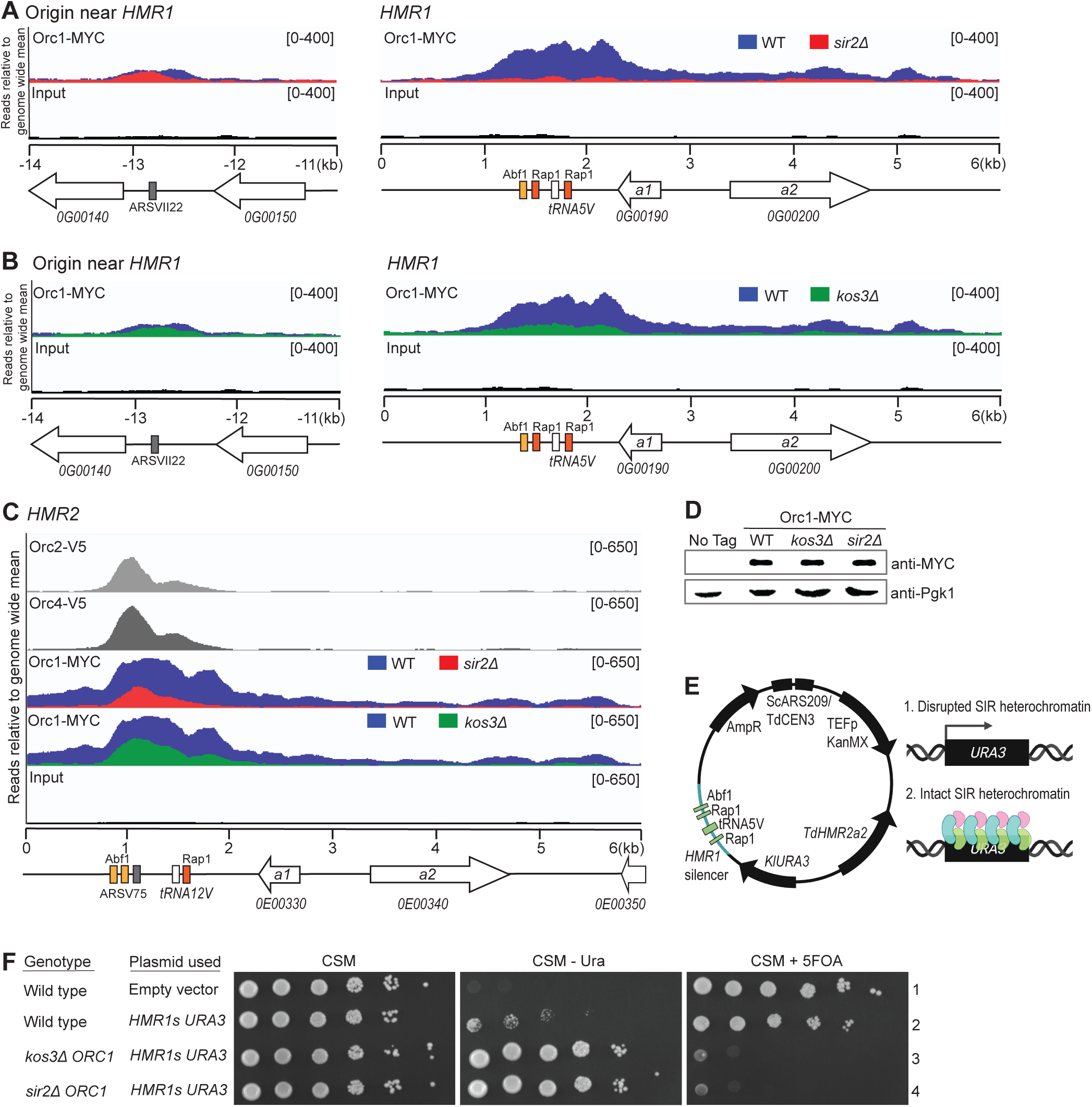
TdSir2 and TdKos3 facilitated the recruitment and spreading of TdOrc1 at *HMR1* and *HMR2*. **(A, B)** Distribution of Orc1-Myc at *HMR1* (right) and the nearest ORC binding site (ARS-VII22; left) in wild-type (LRY3204, blue), *sir2Δ* (LRY3206, red) and *kos3Δ* (LRY3205, green) strains. The ORC binding site is ∼14 kb from the Abf1 binding site at *HMR1*. Signal from input chromatin is shown in black. **(C)** Association of Orc1-Myc at *HMR2* in *sir2Δ* (red) and *kos3Δ* (green) backgrounds and distribution of Orc2-V5, and Orc4-V5 at *HMR2* (Maria *et al*. 2021). **(D)** Immunoblot of Orc1-Myc in wild-type (LRY3204), *kos3Δ* (LRY3205) and *sir2Δ* (LRY3206) backgrounds. The parent strain (JRY10156) is a no tag control, and endogenous 3-phosphoglycerate kinase (Pgk1) is a loading control. **(E)** Diagram of plasmid (pLR1384) used to evaluate transcriptional silencing. The plasmid consists of ∼5 kb fragment of *HMR2* with the **a**1 open reading frame replaced with Kl*URA3* and *HMR2* silencer replaced with *HMR1* silencer (*HMR1s*). The plasmid bears a kanamycin resistance gene for selection. Diagram was created using BioRender.com. **(F)** Expression of *URA3* integrated in place of ***a****1* and adjacent to *HMR1* silencer (*HMR1s*; pLR1384) was measured in wild-type (JRY10156), *kos3Δ* (LRY3205) and *sir2Δ* (LRY3206) backgrounds by assessing growth on media lacking uracil (CSM-Ura, growth requires *URA3* expression) or with 5-FOA (CSM+5FOA, growth requires *URA3* silencing). Empty vector (pRS41K-TdCEN3) containing no *URA3* was used as a control. CSM (left panel) indicates overall growth and equivalent dilutions. All media were supplemented with geneticin to select for plasmids.

Although in *S. cerevisiae* ScSir3 does not require ScSir1 for association with the mating-type loci (Hoppe *et al*. 2002; Rusche *et al*. 2002), the situation may differ in *T. delbrueckii* because the Sir1 ortholog TdKos3 is essential for transcriptional silencing at *HMR2* (Ellahi and Rine 2016). To test if association of TdOrc1 depends on the presence of TdKos3 we conducted ChIP-Seq on Orc1-Myc in a *kos3Δ* background. At *HMR1*, Orc1 association was substantially reduced in the absence of Kos3 (Figure 2B, right panel) but unaltered at the neighboring origin (Figure 2B, left panel). Similarly, at *HMR2* Orc1 was reduced in the absence of Kos3, although it remained associated with the ORC binding site (Figure 2C, track 4). Therefore, unlike ScSir1, TdKos3 is important for the recruitment and spreading of Orc1 across *HMR1* and *HMR2*.

Taken together, these results indicate that both Sir2 and Kos3 are required for TdOrc1 recruitment to *HMR1* as well as its spreading across *HMR2*. Thus, the behavior of TdOrc1 is more like ScSir3, which requires ScSir2 and ScSir4 for spreading, than like ScOrc1, which associates with DNA independently.

### TdSir2 and TdKos3 were required for transcriptional silencing at *HMR1*

The requirement of Sir2 and Kos3 for Orc1 association with *HMR1*, suggests that they are also required for transcriptional repression at *HMR1*, as was shown previously for *HMR2* (Ellahi and Rine 2016). To test this expectation, we used an established plasmid-based assay, in which the entire *HMR* locus was cloned onto a plasmid, and the **a**1 open reading frame was replaced with *URA3*. Expression of this *URA3* reporter was assessed by growth of cells on medium lacking uracil, which is required for cells that do not express *URA3*, and by growth on medium containing 5-FOA, which is toxic to cells expressing *URA3*. We modified the previous plasmid by replacing the *HMR2* silencer with the corresponding region from *HMR1* (Figure 2E). It was not necessary to swap the remainder of the *HMR* locus, because the sequence is nearly identical between *HMR1* and *HMR2* (3939 identities over 3978 aligned nucleotides). Wild-type cells transformed with this plasmid displayed minimal growth in the absence of uracil (Figure 2F, middle panel, row 2) and robust growth in the presence of 5-FOA (Figure 2F, right panel, row 2), indicating silencing of *URA3* within the *HMR* locus. In contrast, *kos3Δ* and *sir2Δ* cells bearing the plasmid displayed robust growth in the absence of uracil (Figure 2F, middle panel, rows 3 and 4) and poor growth on 5-FOA (Figure 2F, right panel, rows 3 and 4), indicating de-repression of the silent locus. A control empty vector with no *HMR2* or *URA3* did not enable growth on plates lacking uracil (Figure 2F, middle panel, row 1) but permitted growth on plates with 5-FOA (Figure 2F, right panel, row 1). Together, these data indicate that both Sir2 and Kos3 are required for repressing *HMR1*.

### The nucleosome-binding BAH domain of TdOrc1 was required for its association and spreading at *HMR1* and *HMR2*

The BAH (bromo adjacent homology) domain of ScSir3 binds nucleosomes, and this ability is critical for the propagation of Sir proteins across silent loci (Onishi *et al*. 2007; Armache *et al*. 2011). To investigate if the BAH domain of TdOrc1 also contributes to silencing, we constructed two mutant alleles of Td*ORC1* designed to abrogate nucleosome binding. First, we deleted the entire bromo adjacent homology (BAH) domain (A2-V213). Second, we replaced proline 179 with alanine (Figure 3A). In ScSir3, a P179A substitution disrupts nucleosome binding and spreading of ScSir3 (Buchberger *et al*. 2008). This is presumably because the main-chain carbonyl of P179 makes an electrostatic interaction with histidine 18 of histone H4 and van der Waals interactions with other residues on the histone H4 tail (Armache *et al*. 2011). Both mutant Orc1 proteins were expressed at levels similar to the wild-type protein (Figure 3B), and neither mutation affected cell viability or growth (data not shown). Consistent with the behavior of ScSir3, the *orc1*-*bahΔ* (Figure 3C, right panel) and *P179A* (Figure 3D, right panel) alleles both disrupted the association of TdOrc1 with *HMR1*, although both mutant proteins retained association with the nearby ORC-binding site (Figure 3C-D, left panels).

**Figure 3:**
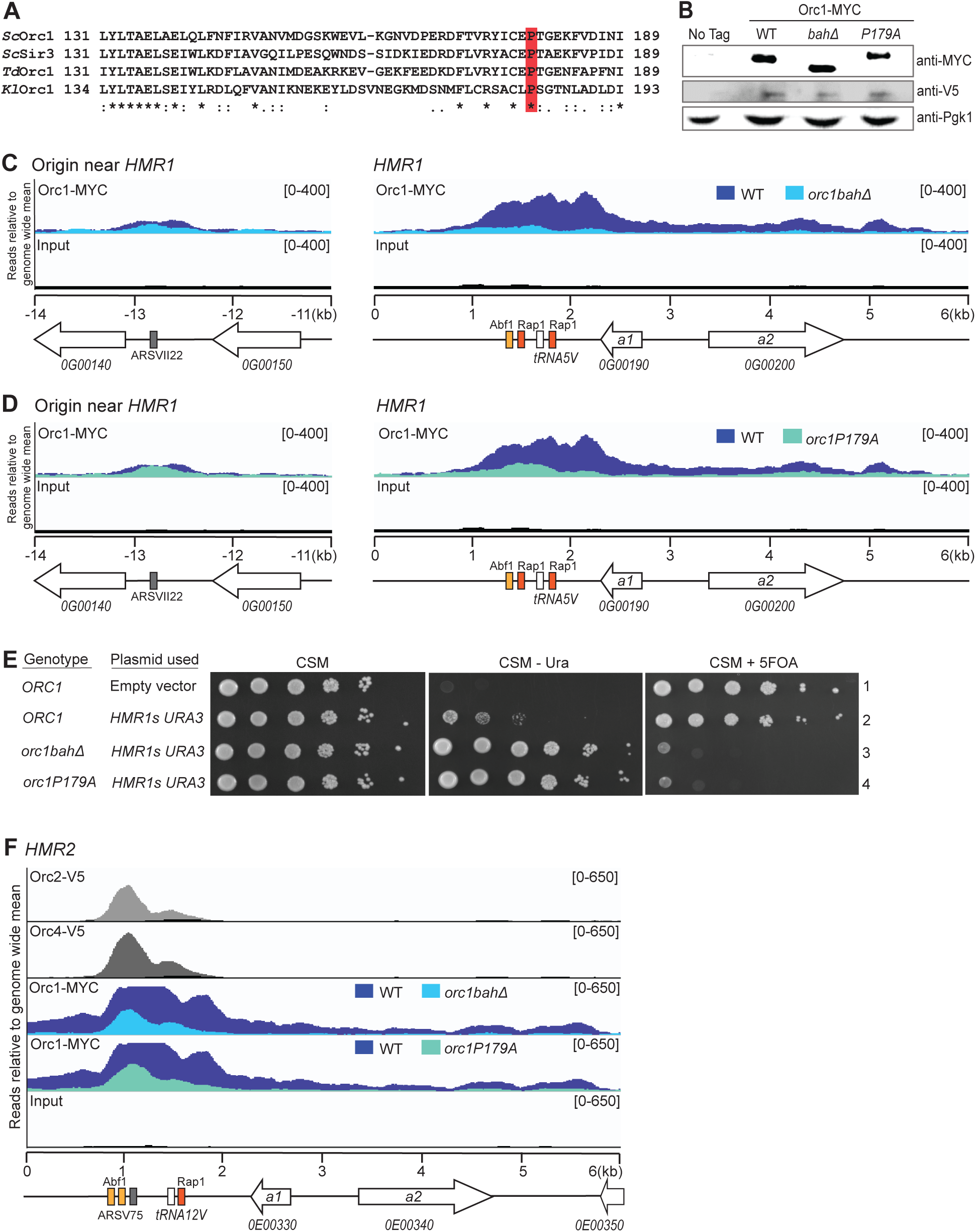
The BAH domain facilitated the recruitment and spreading of TdOrc1 at *HMR1* and *HMR2*. **(A)** Alignment of *T. delbrueckii* Orc1, *S. cerevisiae* Orc1 and Sir3, and *K. lactis* Orc1 showing conservation of the Proline (P) that was mutated (red). **(B)** Immunoblot of wild-type (LRY3211), *bahΔ* (LRY3212), and *P179A* (LRY3215) Orc1-Myc. The anti-V5 band shows the simultaneous expression of Kos3-V5. The parent strain (JRY10156) is a no tag control, and endogenous 3-phosphoglycerate kinase (Pgk1) is a loading control. (**C-D**) Distribution of wild-type Orc1-Myc (LRY3211, blue), *orc1-bahΔ-*Myc (**C**, LRY3212, light blue), and *orc1-P179A*-Myc (**D**, LRY3215, teal) at *HMR1* (right) and the nearest ORC binding site to *HMR1*, ARS-VII22 (left). **(E)** Expression of *URA3* integrated in place of ***a****1* and adjacent to *HMR1* silencer (*HMR1s*; pLR1384) was measured in wild-type *ORC1* (LRY3211), *orc1-bahΔ* (LRY3212), and *orc1-P179A* (LRY3215) backgrounds by assessing growth on media lacking uracil (CSM-Ura, growth requires *URA3* expression) or with 5-FOA (CSM+5FOA, growth requires *URA3* silencing). Empty vector (pRS41K) containing no *URA3* was used as a control. CSM (panel one) was used to assess overall growth and equivalent dilutions. All growth media were supplemented with geneticin to select for plasmids. **(F)** Distribution at *HMR2* of wild-type Orc1-Myc (LRY3211, blue), *orc1-bahΔ-*Myc (LRY3212, light blue), *orc1-P179A*-Myc (LRY3215, teal), Orc2-V5, and Orc4-V5 (Maria *et al*. 2021).

The presence of Orc1 at *HMR1* suggests that it is required for transcriptional repression. To test this idea, we used the plasmid-based expression assay with the *HMR1* silencer and *URA3* integrated in place of ***a****1*. Cells bearing this reporter plasmid and expressing the *orc1*-*bahΔ* or *orc1-P179A* alleles grew robustly in medium lacking uracil (Figure 3E, middle panel, rows 3 and 4) but poorly on 5-FOA (Figure 3E, right panel, rows 3 and 4). In contrast, in a wild-type background there was only weak growth in the absence of uracil and robust growth in the presence of 5-FOA (Figure 3E, row 2). These results indicate that TdOrc1 is required for silencing *HMR1*.

Because the *orc1*-*bahΔ* and *orc1-P179A* alleles disrupt recruitment of Orc1 to *HMR1*, it was not possible to assess whether these alleles also reduce spreading across the locus. However, at *HMR2* Orc1 is recruited as part of ORC, and therefore we were able to assess spreading. In both *bahΔ* and *P179A* strains, Orc1 associated with the ORC binding site within the *HMR2* silencer (Figure 3F). However, spreading across the *HMR2* locus was substantially reduced for *orc1*-*bahΔ* (Figure 3F, track 3, light blue) and *orc1-P179A* (Figure 3F, track 4, teal). These findings suggest that the nucleosome-binding BAH domain is important for spreading of Orc1 across a silenced region, but that it is less important for ORC binding to origins.

To determine if loss of nucleosome binding by TdOrc1 affects association of other Sir proteins with the silent mating-type loci, we investigated the distribution of Kos3-V5 in the same *orc1-bahΔ* and *P179A* cells. Indeed, these *orc1* alleles reduced Kos3 association at both *HMR1* (Figure 4A) and *HMR2* (Figure 4B).

**Figure 4:**
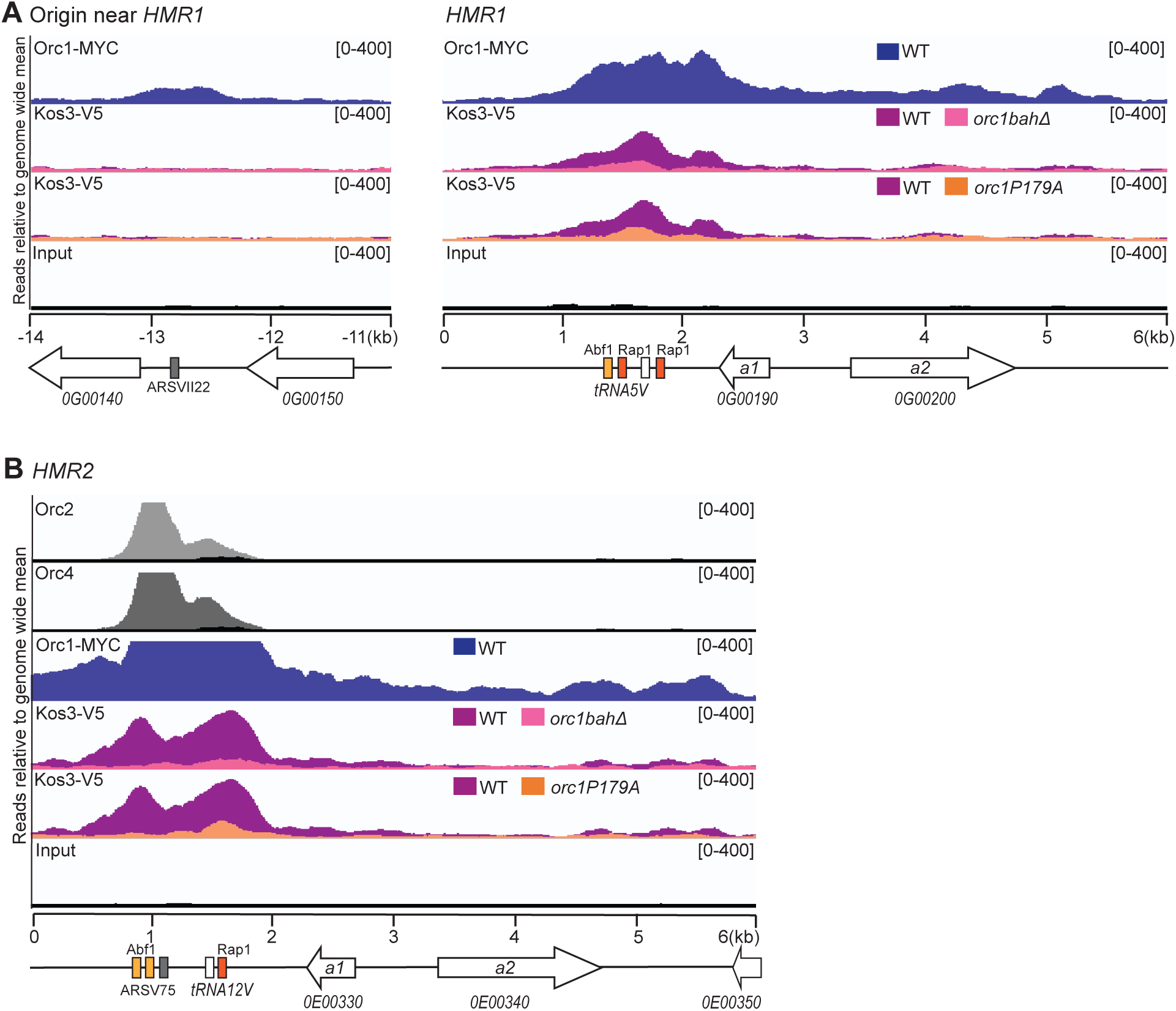
TdOrc1 BAH domain was important for TdKos3 association with *HMR1* and *HMR2*. Distribution of Kos3-V5 association in wild-type (LRY3211, purple), *orc1-bahΔ-*Myc (LRY3212, pink) and *orc1P179A*-Myc (LRY3215, orange) strains at *HMR1* **(A)** and *HMR2* **(B)**. Distribution of Orc1-Myc (blue), Orc2-V5 (light grey) and Orc4-V5 (dark grey) are shown for comparison (Maria *et al*. 2021). Signal from input chromatin is shown in black.

### An ORC binding site was not required for Orc1 association with *HMR1*

In *S. cerevisiae*, Orc1 contributes to silencing by recruiting Sir1, which in turn recruits other Sir proteins to the silent locus (Fox *et al*. 1997; Gardner *et al*. 1999). In this role, ScOrc1 acts as part of the ORC complex, binding to silencers flanking the silent mating-type loci (Foss *et al*. 1993). A similar function for TdOrc1 seems unlikely at *HMR1* given the lack of Orc2 and Orc4 enrichment (Figure 1D). Nevertheless, we examined whether the nearest ORC binding site (∼14 kb away) recruits Sir proteins to *HMR1*. We deleted this ORC binding site, ARS-VII22, in a strain lacking *HMR2* so that the ChIP signal would be derived solely from *HMR1*. To confirm the disruption of ORC binding at this locus, we immunoprecipitated Orc1-Myc and analyzed the co-precipitated DNA by qPCR. Deleting ARS-VII22 abolished Orc1 binding at this locus, as assessed using primers adjacent to the deleted region (Figure 5B, left panel). However, deleting this ORC binding site did not decrease binding at another control origin, ARS-V931 (Figure 5A). Importantly, Orc1 remained associated with *HMR1* in *arsvii22Δ* cells (Figure 5B, right panel). Therefore, Orc1 association at *HMR1* was not dependent on ORC binding at ARS-VII22. This finding, combined with the lack of ORC binding sites within *HMR1* (Figure 1), indicates that TdOrc1 is not a silencer binding protein at *HMR1*.

**Figure 5:**
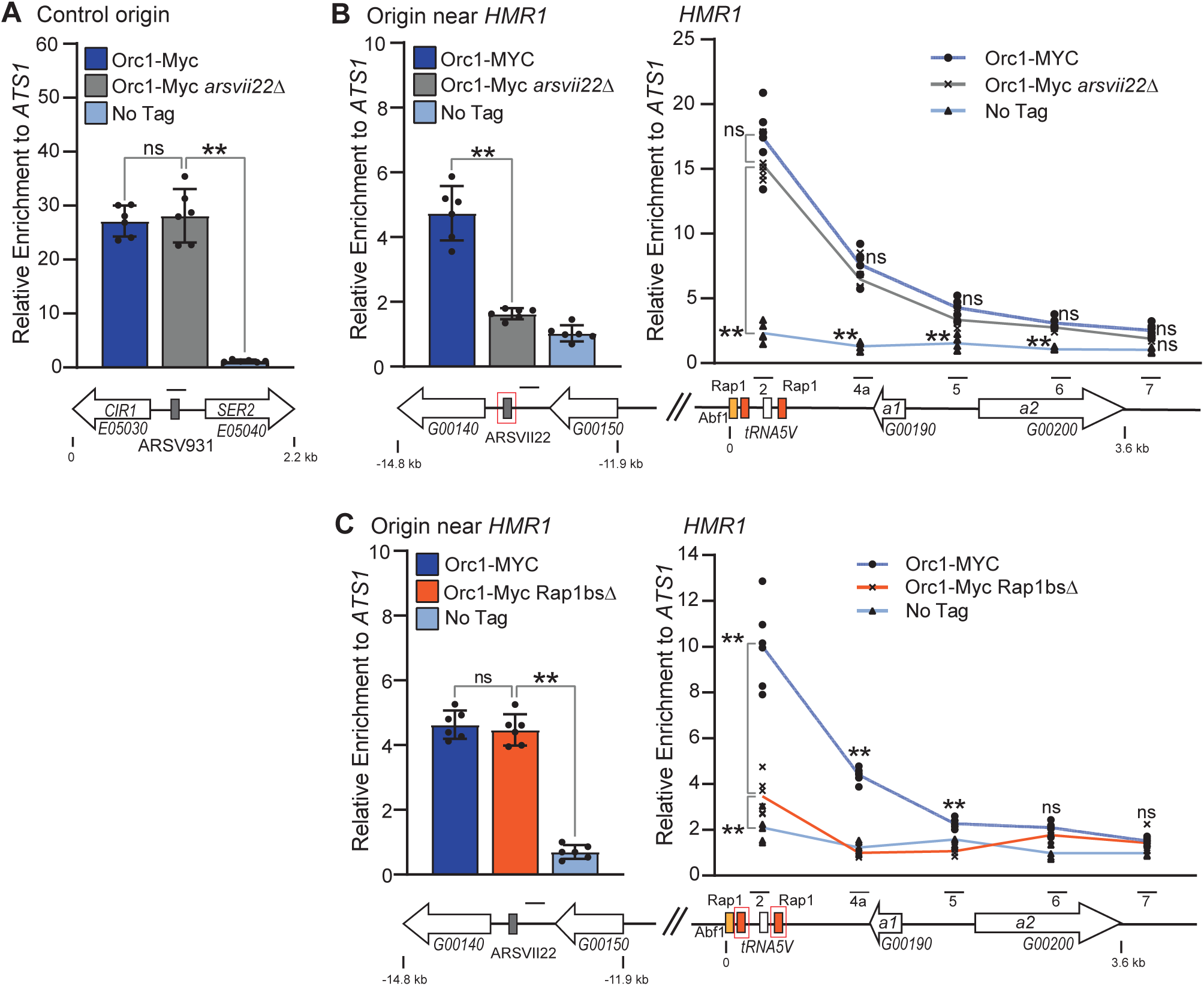
Binding sites for Rap1, but not ORC, were required for Orc1 association with *HMR1*. **(A)** TdOrc1 enrichment at a control origin, ARS-V931, was determined by chromatin IP followed by quantitative PCR (ChIP-qPCR) in strains with (dark blue, LRY3208 and 3209) and without (grey, LRY3334 and 3335) the ORC binding site nearest to *HMR1*, ARS-VII22. **(B)** TdOrc1 enrichment at *HMR1* (right) and the nearest ORC-binding site (ARS-VII22, left) in the presence (dark blue) and absence (grey) of ARS-VII22. **(C)** TdOrc1 enrichment at ARS-VII22 (left) and *HMR1* (right) in strains with wild-type *HMR1* (dark blue, LRY3208 and 3209) or lacking the Rap1 <u>b</u>inding <u>s</u>ites (*rap1<u>bs</u>Δ*, red, LRY3406 and 3407). The strains used for these experiments lacked *HMR2*. Mock immunoprecipitation using anti-Myc antibody and the untagged parent strain (JRY10156) is shown in light blue. The y-axis represents the relative enrichment to the control locus *ATS1*. Solid black lines represent the position of the qPCR primers. The bars represent the average of six independent IPs with error bars representing SEM. Asterisks show statistically significant differences (\*\**p* < 0.005, \**p* < 0.01, ns = *p* > 0.05) determined using two-tailed Student’s t-test.

### Rap1 binding sites were required for Orc1 association with *HMR1*

For comparison with the ORC binding site deletion, we deleted the binding sites for Rap1, another protein known to bind silencers in *S. cerevisiae* (Moretti *et al*. 1994). Rap1 binding sites are essential for establishing silencing at *HMR2* in *T. delbrueckii* (Ellahi and Rine 2016) and three of four silencers in *S. cerevisiae* (Shore and Nasmyth 1987; Buchman *et al*. 1988). We therefore deleted the two putative Rap1 binding sites at *HMR1* (Figure 5C, panel two, red boxes) in a strain lacking *HMR2*. We then immunoprecipitated Orc1-Myc and analyzed the DNA by qPCR in both wild-type (Figure 5C, right panel, dark blue) and *rap1*_*bs*_*Δ* (Figure 5C, right panel, grey) backgrounds. Deleting Rap1 binding sites significantly reduced Orc1 association at *HMR1* compared to wild-type cells, while Orc1 enrichment at the neighboring ORC binding site, ARS-VII22, was unaffected (Figure 5C, left panel, grey and dark blue). These results indicate that the Rap1 binding sites, but not the nearby ORC binding site, are required for Orc1 recruitment to *HMR1*.

### An ORC binding site at *HMR2* was dispensable for transcriptional silencing

Unlike *HMR1*, there is an ORC binding site in the silencer of *HMR2* (Figure 1E). Therefore, it was possible that TdOrc1 acts as a silencer-binding protein at this locus. To investigate this possibility, we adapted the plasmid based *URA3* reporter assay (Ellahi and Rine 2016). Starting with a plasmid containing the full-length *HMR2* locus, we deleted the 17 bp conserved ORC binding motif (Maria *et al*. 2021) in the silencer (Figure 6A). To assess whether this deletion (*arsV75Δ*) reduced ORC binding, we conducted chromatin immunoprecipitation on Orc2, which is part of ORC but not involved in silencing. We observed reduced Orc2 association near the ORC binding site for the *arsV75Δ* plasmid compared to a plasmid with an intact silencer (Figure 6B, right panel, red box). However, there was residual Orc2 association that we attributed to the plasmid origin (*ScARS209*), ∼3.4 kb away. Indeed, Orc2 was associated with *ScARS209* on plasmids with and without the ORC binding site at the silencer (Figure 6B, left panel). As a second way to assess whether the *arsV75Δ* deletion eliminated ORC binding, we also deleted the plasmid origin. We found that when *T. delbrueckii* cells were transformed with a plasmid lacking both *ScARS209* and the ORC binding motif in the silencer, only tiny yeast colonies were obtained, consistent with the plasmid not being replicated. In contrast, when cells were transformed with a plasmid that had the silencer ORC binding motif but lacked *ScARS209*, the normal sized colonies were observed (Figure 6C). Therefore, the silencer ORC binding site can serve as an origin, and this capacity is lost in the *arsV75Δ* plasmid. Thus, the *arsV75Δ* deletion successfully eliminated ORC binding.

**Figure 6:**
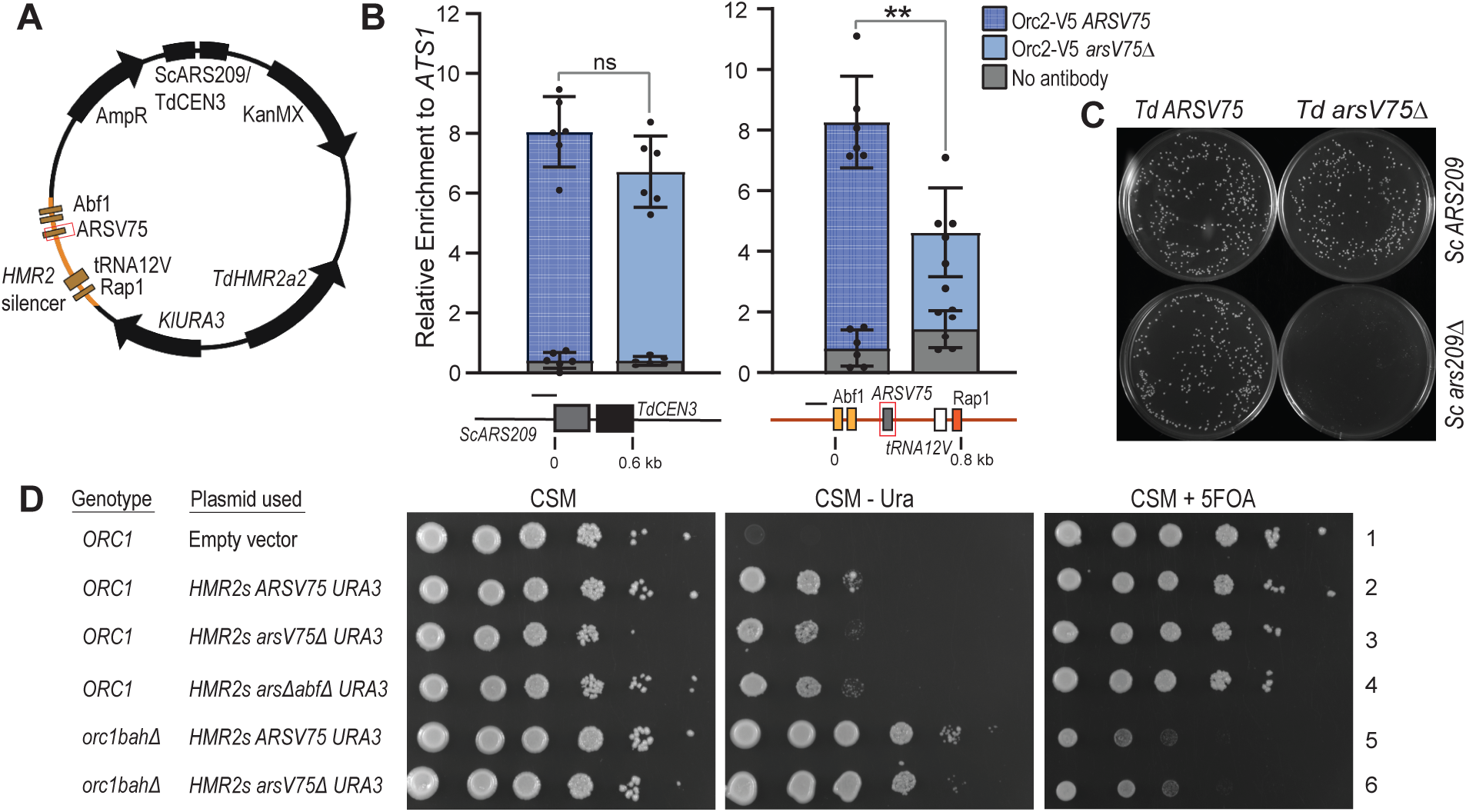
An ORC binding site at *HMR2* was dispensable for transcriptional silencing. **(A)** Diagram of reporter plasmid to evaluate role of Orc1 binding at the *HMR2* silencer in gene expression. The plasmid carries a ∼5 kb fragment of *HMR2* with the **a**1 open reading frame replaced with Kl*URA3*. The plasmid bears a Kanamycin resistance gene for selection in geneticin. Diagram was created using BioRender.com. **(B)** ChIP-qPCR showing TdOrc2-V5 enrichment at the plasmid origin, *ScARS209* (left) or the *HMR2-*associated ORC binding site, *TdARS-V75* (right), in the presence (dark blue, pLR1377) or absence (light blue, pLR1382) of the ORC binding site. An Orc2-V5 strain (LRY3142) was transformed with both plasmids. No antibody controls (grey) are overlayed on respective IP signals. The y-axis represents the relative enrichment to the control locus *ATS1*. The bars represent the average of six independent IPs with error bars representing SEM. Asterisks show statistically significant differences (\*\**p* < 0.005, \**p* < 0.01, ns = *p* > 0.05) determined using two-tailed Student’s t-test. Black lines in the schematic represent the position of qPCR primers. **(C)** Plasmid stability assay to assess contribution of ORC binding sites. Wild-type *T. delbrueckii* cells (JRY10156) were transformed with plasmids with both the plasmid and silencer ORC binding sites (pLR1377), *arsV75Δ* (pLR1382), *ars209Δ* (pLR1401), or both *arsV75Δ* and *ars209Δ* (pLR1400). Transformants were selected on geneticin and imaged after two days. **(D)** Expression of *URA3* integrated in place of ***a****1* and adjacent to *HMR2* silencer in wild-type *ORC1* (JRY10156) or *orc1-bahΔ* (LRY3212) strains. Yeast were transformed with plasmids containing no *URA3* (empty vector, pRS41K-TdCEN3, row 1), *HMR2****a****1::URA3* with wild-type silencer (pLR1377, rows 2 and 5), no ORC binding site in the silencer (*arsV75Δ*, pLR1382, rows 3 and 6), or no ORC or Abf1 binding sites (*arsΔ+abf1*_*bs*_*Δ*, pLR1399, row 4). Cells were grown on CSM (left) to assess growth and dilution, CSM-Ura (middle panel, growth requires *URA3* expression), and CSM+5FOA (right panel, growth requires *URA3* silencing). All plates were supplemented with geneticin to select for the plasmids.

Next, we assessed *URA3* silencing on the *arsV75Δ* plasmid. Cells containing this plasmid grew similarly to those containing a plasmid with the wild-type silencer; they grew poorly on CSM-Ura (Figure 6C, middle panel, rows 2 and 3) and well on 5-FOA (Figure 6C, right panel, rows 2 and 3). This result indicates that loss of the ORC binding site did not affect the silencing of *URA3* and therefore that an ORC binding site at *HMR2* is not required to establish silencing. However, it remained possible that there is redundancy in the silencer and that the ORC binding site is not needed when other silencer binding proteins are present. In fact, for the *S. cerevisiae HMR-E* silencer, two of three binding sites must be deleted to produce a loss of silencing (Brand *et al*. 1987). We therefore created a larger deletion in which the ORC binding site and both Abf1 binding sites were deleted. Cells containing this plasmid also grew poorly on CSM-Ura (Figure 6C, middle panel, row 4) and well on 5-FOA (Figure 6C, right panel, row 4), indicating that even loss of the ORC and Abf1 binding sites did not derepress *URA3*. A similar result was previously observed (Ellahi and Rine 2016). Therefore, the ORC binding site in the *HMR2* silencer is not important for silencing. In contrast, Orc1 itself is required to silence *HMR2*; in an *orc1-bahΔ* strain *URA3* is expressed from the plasmid with the wild-type silencer (Figure 6C, row 5). Together, these results indicate that TdOrc1 is important at *HMR2* but not as a silencer-binding protein.

### Some Kos3 proteins from *Torulaspora* species lack an ORC interacting region (OIR)

Two of our findings suggest that for TdKos3, binding to Orc1 at the silencer is not its primary role in silencing. First, TdKos3 has the same distribution at mating-type loci as TdSir2, TdSir4, and TdOrc1, rather than being restricted to an ORC binding site (Figure 1D-F). Second, ORC does not act as a silencer-binding protein at *HMR1* (Figure 5) or *HMR2* (Figure 6). To explore whether an interaction between TdKos3 and TdOrc1 is likely, we compared the ORC interacting regions (OIR) of Sir1 and Kos3 proteins from species across the *Saccharomycetaceae* family (Figure 7; see Figure 1A for species relationships). To identify the OIR, each protein was submitted to NCBI blastp (https://blast.ncbi.nlm.nih.gov/Blast.cgi), which returned the boundaries of a pre-defined domain that corresponds to the OIR (called “Sir1 superfamily,” architecture ID 11189688). Interestingly, for several Kos3 proteins from the *Torulaspora* and *Zygotorulaspora* genera, including *T. delbrueckii*, this “Sir1 superfamily” domain was not identified by the blastp analysis (Figure 7), suggesting the OIR is absent or highly diverged from the consensus. Nevertheless, other conserved sequences that are shared among Kos3/Sir1 proteins could be identified in these proteins using the motif finder MEME (Bailey *et al*. 2009), demonstrating that these proteins are Kos3 homologs. We also found that the Kos3 proteins lacking an OIR aligned poorly to other Kos3 proteins and had lower sequence similarity. Therefore, unlike ScSir1, TdKos3 does not appear to have an OIR that would enable it to bind to the BAH domain of TdOrc1.

**Figure 7:**
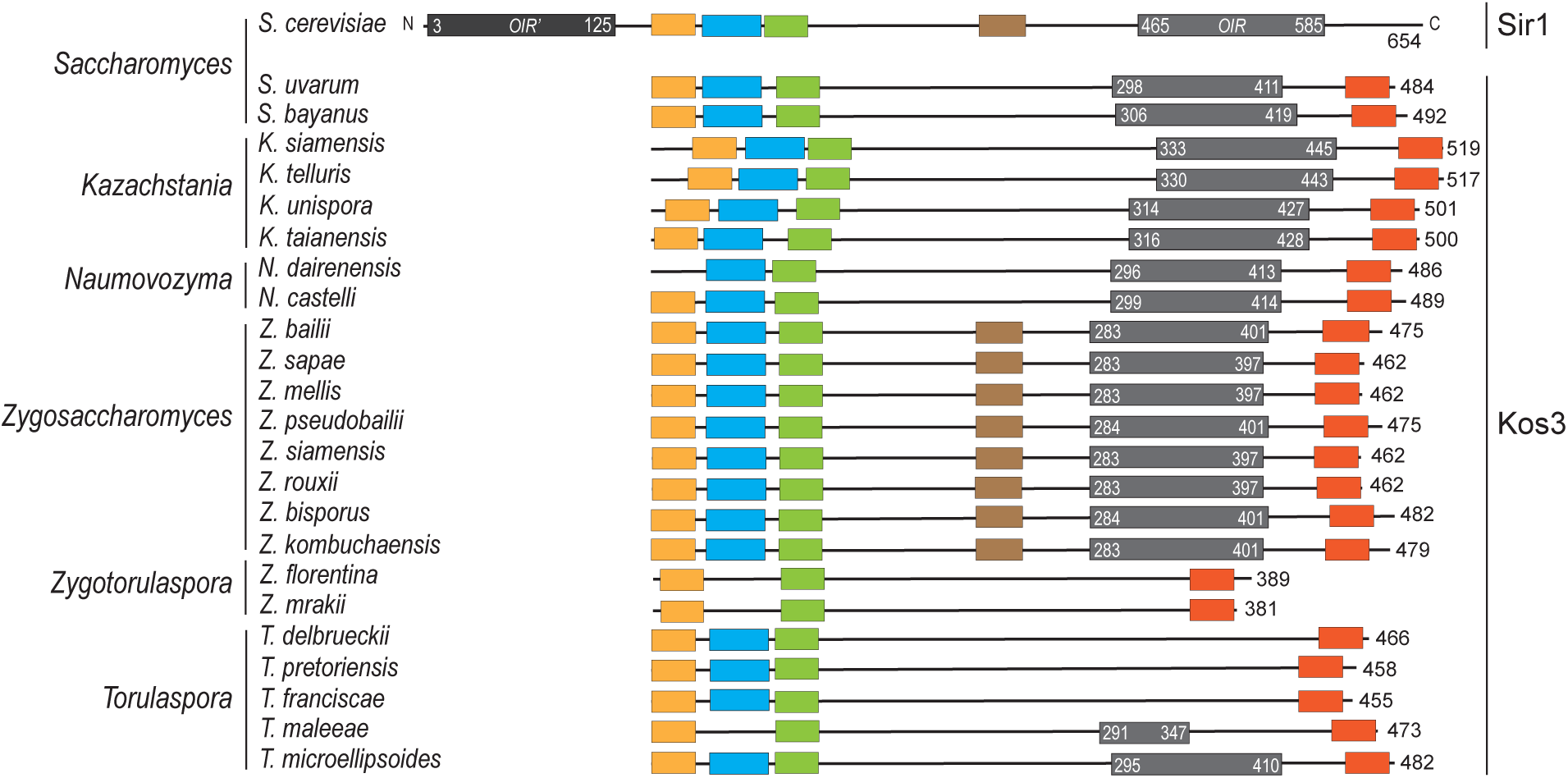
Some Kos3 orthologs lack an Orc Interacting Region (OIR). Sequences of Kos3 orthologs were obtained from NCBI, and analyzed for the presence of conserved domains. OIR domains (light grey boxes) were identified by submitting each protein as a query to NCBI blastp. and the program returned the boundaries of the defined “Sir1 superfamily” domain that corresponds to the OIR. Other conserved regions (colored boxes) were identified using the MEME suite (Bailey *et al*. 2009) to search for sequence motifs present zero or one time in each member of the protein set. Lengths of the proteins are relative to their sizes. *S. cerevisiae* Sir1 is included for comparison. ScSir1 has a C-terminal OIR (light grey) and N-terminal OIR’ (dark grey).

### Silencers at the mating-type loci were associated with TdRap1 and TdAbf1 but not TdSum1, TdReb1, or TdUme6

Given that ORC does not serve as a silencer-binding protein in *T. delbrueckii*, we were interested to identify other proteins that do. In *T. delbrueckii* only Rap1 bindings sites are documented to be required for silencing ((Ellahi and Rine 2016) and Figure 5). Whereas, in *S. cerevisiae*, three silencer-binding proteins are known: ORC, Rap1, and Abf1. The other species in which silencer-binding proteins have been studied is *K. lactis*, which uses different DNA-binding proteins to initiate silencing, namely Reb1, Ume6 and Sum1 (Sjostrand *et al*. 2002; Hickman and Rusche 2009; Barsoum *et al*. 2010). To elucidate which proteins associate with *T. delbrueckii* silencers, we tagged the five DNA-binding proteins that act at silencers in *S. cerevisiae* or *K. lactis* and then conducted chromatin IP to determine whether these proteins are present at predicted silencers. Silencers coincide with the regions of highest Sir protein association and are defined at *HML, HMR2* (Ellahi and Rine 2016) and *HMR1* (Figures 1 and 5).

TdRap1 and TdAbf1 were both enriched at *HMR1* (Figure 8B) and *HMR2* (Figure 8C), whereas only TdRap1 was enriched at *HML* (Figure 8D). For the other DNA-binding proteins, we did not observe significant enrichment at the silencers, although the immunoprecipitation was successful based on enrichment at control loci: TdUme6 at *SPO11*, TdReb1 at *RDN25-1* and TdSum1 at *CDA2* (Figure 8E). To better define the silencers, we identified putative binding sites Rap1 and Abf1. For TdRap1, we determined a consensus binding motif using ChIP-Seq data (unpublished) and located matches to this motif that correspond to peaks of Rap1 enrichment as assessed by ChIP-Seq (unpublished) and ChIP-qPCR (Figure 8). Moreover, deletion of these motifs at *HMR1* lead to loss of Orc1 association (Figure 5C). We also identified putative binding sites for Abf1 as matches to the known motif for *S. cerevisiae* Abf1. At *HMR1* and *HMR2* these putative binding sites correspond to the locations of TdAbf1 enrichment (Figures 8B and C). Overall, these results indicate that TdRap1 and TdAbf1 are present at the silencers of *T. delbrueckii* mating-type loci.

**Figure 8:**
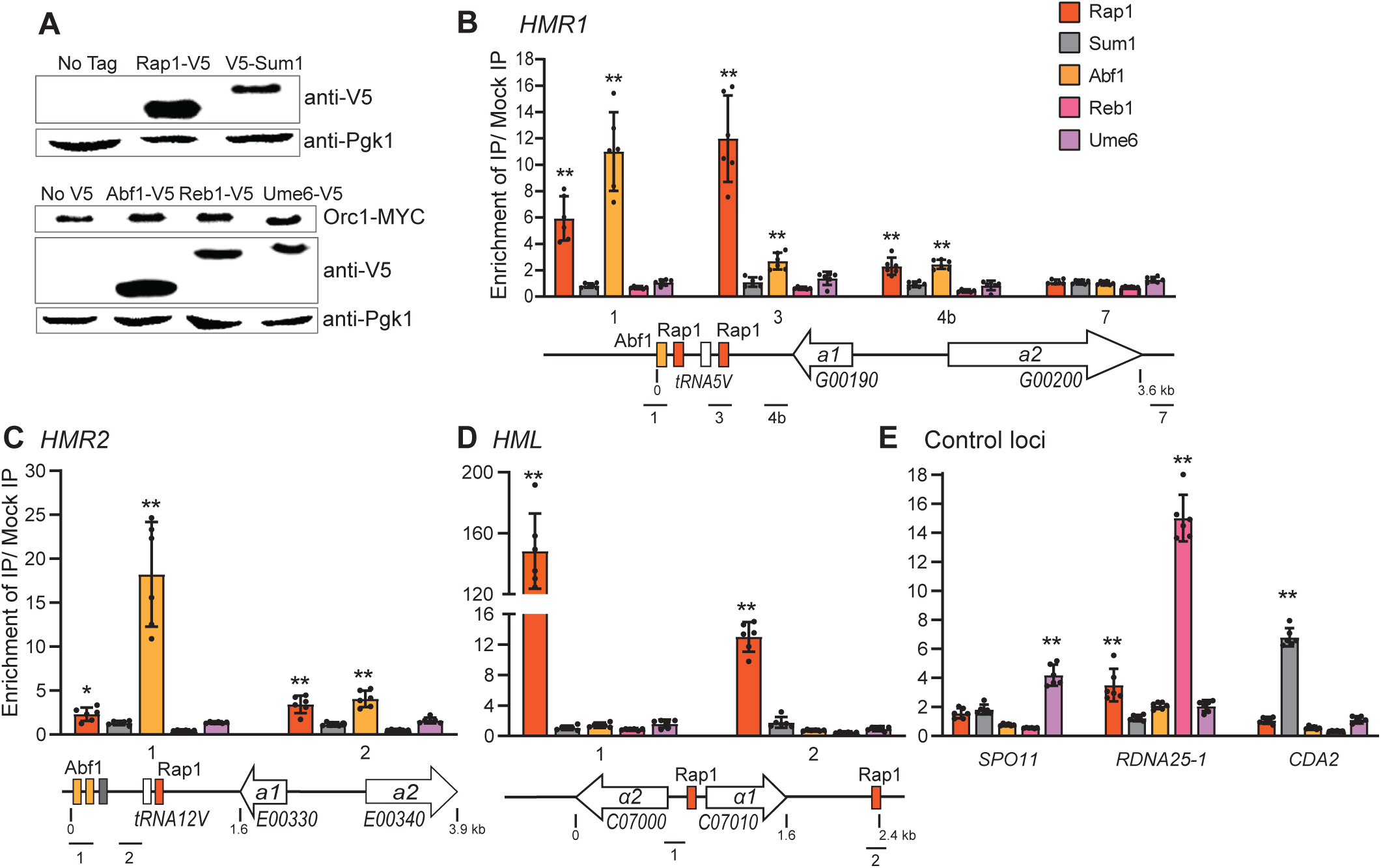
Rap1 and Abf1 associated with the silencers at the *T. delbrueckii* mating-type loci. **(A)** Immunoblot of V5 tagged Rap1 (JRY10207), Sum1 (LRY3172), Abf1 (LRY3336), Reb1 (LRY3338) and Ume6 (LRY3340) proteins. The parent strains (JRY10156 and LRY3208) were included as no V5 tag controls. Endogenous 3-phosphoglycerate kinase (Pgk1) was detected as a loading control. **(B-D)** Association of the silencer binding candidates, Rap1 (orange, JRY10207) Abf1 (yellow, LRY3336 and LRY3337), Sum1 (grey, LRY3172 and 3173), Reb1 (blue, LRY3338 and 3339), and Ume6 (green, LRY3340 and 3341) at *HMR1* **(B)**, *HMR2* **(C)**, and *HML* **(D)** loci. Solid black lines represent the positions of the qPCR primers. **(E)** Association of Sum1 (grey), Reb1 (blue) and Ume6 (green) with other genomic loci to confirm successful immunoprecipitation. The y-axis represents ratio of IP over mock IP: IP enrichments were first calculated relative to the control locus *ATS1* and then normalized to the mock IP Mock IPs were anti-Myc and anti-V5 antibody pulldowns in untagged parent strains (JRY10156 and LRY3208). The bars represent the average of six independent IPs with error bars representing SEM. Asterisks show statistically significant differences (\*\**p* < 0.005, \**p* < 0.01, ns = *p* > 0.05) when IP sample was compared to mock IP using two-tailed Student’s t-test.

## DISCUSSION

In this study, we examined the hypothesis that the appearance of a Sir1-like protein prior to the whole genome duplication enabled Orc1 to become a silencer-binding protein. To do so, we characterized the role of Orc1 in heterochromatin formation in the non-duplicated yeast species *T. delbrueckii*, which belongs to the earliest clade of yeast to encode a Sir1-like protein, Kos3. However, we did not observe a silencer binding function for TdOrc1. Nevertheless, we did find that TdOrc1 contributed to heterochromatin formation through its nucleosome-binding BAH domain, much as ScSir3 and KlOrc1 do. In keeping with TdOrc1 not serving as a silencer-binding protein, we extended previous observations (Ellahi and Rine 2016) that TdKos3 plays a different role in heterochromatin formation than ScSir1 does. Namely, TdKos3 and other Sir proteins depend on one another to spread across the silenced loci. Thus, the protein composition of the heterochromatin in *T. delbrueckii* is distinct compared to other species that have been examined (Figure 9).

**Figure 9:**
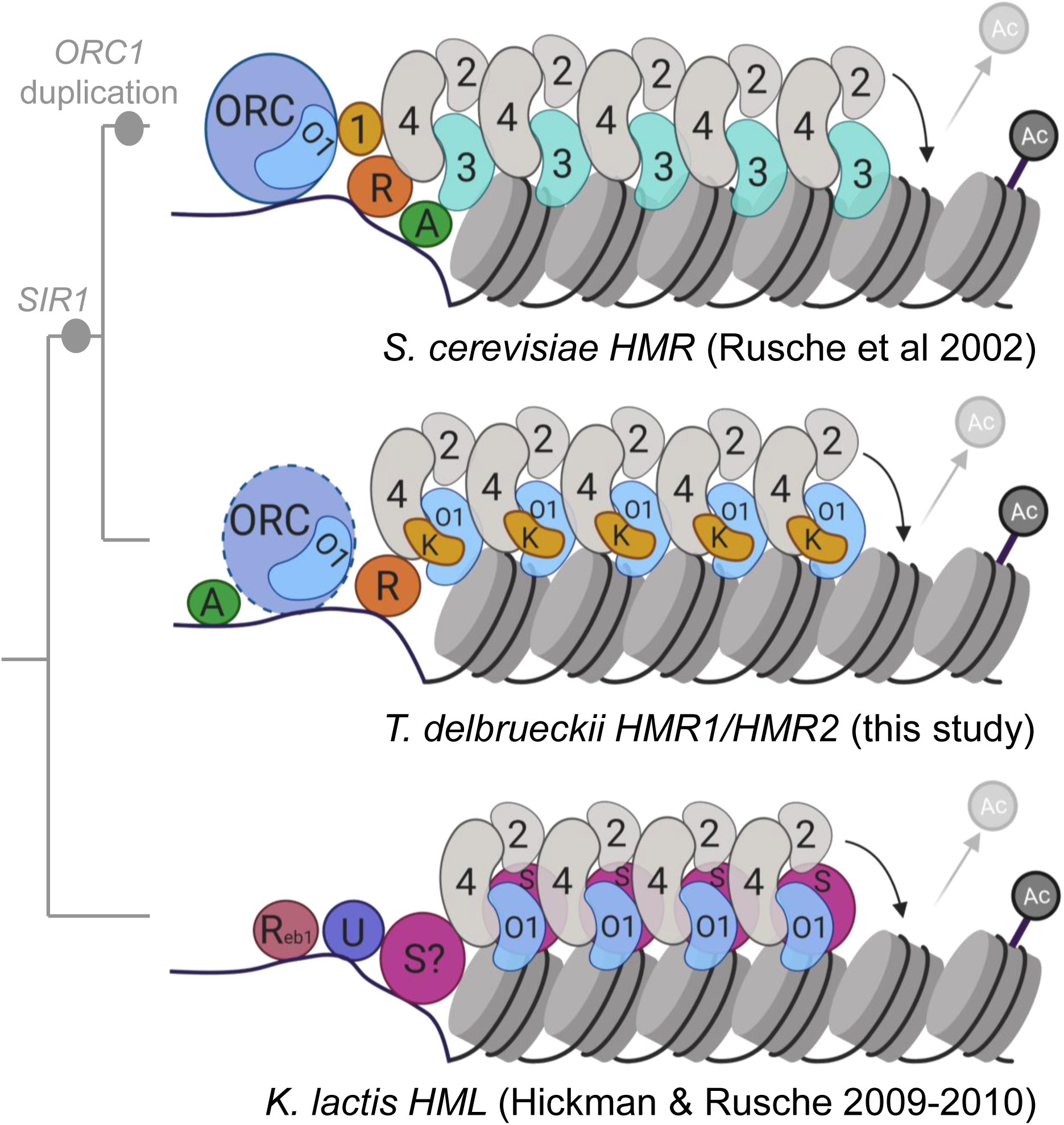
Composition of SIR heterochromatin in three budding yeast. Model for SIR heterochromatin at *S. cerevisiae HMR* (top), *T. delbrueckii HMR2* (middle; model based on this study), and *K. lactis HML* (bottom). *ORC1* duplication – duplication resulting in *ORC1* and *SIR3. SIR1* – appearance of *SIR1* family. ORC – Origin Recognition Complex (violet), 1 – Sir1 (yellow), 2 – Sir2 (grey), 4 – Sir4 (grey), 3 – Sir3 (teal), R – Rap1 (brick red), A – Abf1 (green), Ac – acetyl group on H4K16 (grey), O1 – Orc1 (blue), K – Kos3 (yellow), R_eb1_ – Reb1 (coral), U – Ume6 (dark violet), and S – Sum1 (pink). Figure was created using BioRender.com.

Several lines of evidence support the conclusion that TdOrc1 does not behave as a silencer-binding protein. First, there are no ORC binding sites at two of the mating-type loci (*HMR1* and *HML*, Figure 1D and F), and the ORC binding site at *HMR2* was not required for silencing (Figure 6). In addition, TdOrc1 did not associate with the mating-type loci independently of other silencing proteins, but instead required Sir2 and Kos3 (Figure 2). Moreover, recruitment of Orc1 to *HMR1* required the Rap1 binding sites in the silencer but not the nearest ORC binding site (Figure 5). Thus, TdOrc1 does not behave as a silencer-binding protein. We had speculated that the emergence of a Sir1-like protein would coincide with Orc1 acting as a silencer binding protein. However, for *T. delbrueckii*, having a Sir1-like protein is not sufficient for Orc1 to serve as a silencer binding protein.

To better understand the lack of silencer binding function in TdOrc1, we compared the domain structure of TdKos3 to other Kos3 orthologs and found that TdKos3 lacks an ORC interacting region (OIR) (Figure 7). This domain was identified and characterized in *S. cerevisiae* Sir1 to interact with the Orc1 BAH domain (Gardner *et al*. 1999; Hou *et al*. 2005). Thus, without an OIR, TdKos3 is unlikely to bind to the TdOrc1 BAH domain. Given the widespread distribution of the OIR among other Kos3 orthologs and Sir1-like proteins (Figure 7 and (Gallagher *et al*. 2009)), we infer that the ancestral Kos3 had an OIR, which was subsequently lost in the *Torulaspora* lineage. Thus, it remains possible that other non-duplicated Orc1 proteins are silencer-binding proteins.

Despite lacking an OIR, TdKos3 plays an important role in transcriptional silencing. Deletion of *KOS3* derepresses transcription and reduces the association of Sir2, Sir4, and Orc1 with the mating type loci ((Ellahi and Rine 2016) and Figure 2). Moreover, unlike ScSir1, TdKos3 is distributed across the mating-type loci in the same pattern as Sir2, Sir4 (Ellahi and Rine 2016), and Orc1 (Figure 1), and it requires the Orc1 BAH domain for association with these loci (Figure 4). Taken together these findings suggest that Kos3 forms a complex with Sir2, Sir4, and Orc1, and that these proteins depend on one another to assemble and spread across silenced loci (Figure 9). Thus, TdKos3 appears to be an integral part of the silencing complex rather than a bridge between ORC and the other silencing proteins, as ScSir1 is. It remains to be determined whether other Kos3 orthologs behave similarly to TdKos3, and if so whether having an OIR that binds Orc1 facilitates assembly with Sir proteins, bridging to ORC at a silencer, or both. It is notable that the set of heterochromatin proteins that spreads differs in *S. cerevisiae, T. delbrueckii*, and *K. lactis*. In *S. cerevisiae* Sir2, Sir3, and Sir4 but not Sir1 spread away from silencers (Hoppe *et al*. 2002; Rusche *et al*. 2002). In *K. lactis* Sum1 spreads with Sir2, Orc1, and Sir4 (Hickman and Rusche 2009; Hickman and Rusche 2010). We show here that in *T. delbrueckii* Kos3 joins Sir2, Orc1, and Sir4. Hence, we postulate that prior to the emergence of Sir3 through duplication of *ORC1*, the Sir2/Orc1/Sir4 complex was insufficient on its own and required other proteins, such as Kos3 or Sum1, for stability. For example, because ORC is required for DNA replication and viability, Orc1 may not have been able to evolve optimal interactions with other Sir proteins while maintaining surfaces for assembly into the ORC complex. However, after duplication, Sir3 acquired new properties in the AAA+ base subdomain (Hanner and Rusche 2017), which may have strengthened the Sir2/Sir3/Sir4 complex such that there was no longer a need for a protein such as Kos3 or Sum1. Such a scenario could account for TdKos3 being essential for silencing, whereas ScSir1 is not.

Our finding that TdOrc1 promotes the spreading of heterochromatin through its nucleosome-binding BAH domain (Figure 3) reinforces the notion that this property arose prior to silencer binding. We previously found that Orc1 from *K. lactis* also promotes spreading of heterochromatin but does not bind silencers (Hickman and Rusche 2010). It is therefore probable that the common ancestor of *T. delbrueckii* and *K. lactis* also employed Orc1 in this way. Thus, considering this ancient role of the Orc1 BAH domain in promoting the spreading of Sir proteins and that the BAH domain of Orc1 is conserved across eukaryotes, the BAH domain might play a similar role in species beyond yeast. Moreover, ORC binding of HP1 (heterochromatin protein 1) in *Drosophila* and *Xenopus* (Pak *et al*. 1997; Shareef *et al*. 2001) may not be analogous to the ORC-assisted Sir1 binding in *S. cerevisiae*.

Finally, we found that the silencer architecture of *T. delbrueckii* more resembles that of *S. cerevisiae* than *K. lactis. S. cerevisiae* silencers have binding sites for Rap1, Abf1, and ORC, whereas *K. lactis* silencers have binding sites for Reb1, Ume6, and potentially Sum1. All three presumed *T. delbrueckii* silencers have Rap1 binding sites (Figure 8), and Rap1 is critical for silencer function at *HML* and *HMR2* (Ellahi and Rine 2016) as well as *HMR1* (Figure 5). Abf1 binding sites are found at both *HMR* silencers (Figure 8), and ORC binds to the *HMR2* silencer (Figure 1E). However, there is no evidence that either Abf1 or ORC contributes to silencing in *T. delbrueckii*. Thus, their presence at silencers may be vestigial. We did not detect the proteins associated with *K. lactis* silencers at *T. delbrueckii* silencers (Figure 8). The shift in silencer architecture between *K. lactis* and *T. delbrueckii* could indicate that there is no adaptive advantage to having a particular set of silencer binding proteins. On the other hand, the similarity between *S. cerevisiae* and *T. delbrueckii* silencers suggests that there is an evolutionary advantage to maintaining a Rap1-Abf1-ORC type silencer.

In summary, we found that the presence of a Sir1-like protein does not necessarily coincide with Orc1 having a silencer-binding function. Thus, both Orc1 and Sir1/Kos3 originally had different roles in heterochromatin formation than they do now in *S. cerevisiae*, illustrating the flexibility in how these proteins achieve silencing.

## Supporting information

Reagent Table

## ACKNOWLEDGEMENTS

We thank Esther Oladunjoye, E. Ray Santillano, and Jannatul Ferdoes Shoma for technical assistance, Michael Buck and Tao Liu for suggestions on ChIP-Seq analysis, Jasper Rine and Aisha Ellahi for strains and plasmids, Aisha Ellahi for comments on this manuscript, and members of the Rusche lab for suggestions and support. Bioinformatic analysis was supported by the Center for Computational Research at the University at Buffalo (http://hdl.handle.net/10477/79221).

