## Supplemental Methods for "The DNA replication protein Orc1 from the yeast *Torulaspora delbrueckii* is required for heterochromatin formation but not as a silencer-binding protein"

### SUPPLEMENTAL MATERIALS AND METHODS

#### Yeast strain construction

Yeast strains used in this study are listed in the Reagent Table, and the details of their construction are provided in Tables S1 and S2. Yeast strains were derived from JRY10156 (ELLAHI AND RINE 2016), a *Torulaspora delbrueckii MAT*α *ura3Δ0 trp3(G466A)-1* strain descended from NRRL Y-866.

To add epitope tags to proteins of interest, tagged alleles were first generated on plasmids (Table S1). Genes, including ~500 bp flanking the open reading frame, were amplified from genomic DNA (JRY10156) and ligated into pRS414 (SIKORSKI AND HIETER 1989). The genes were *T. delbrueckii ORC1* (*TDEL0D00910*), *KOS3* (*TDEL0E00350*), *ABF1* (*TDEL0A02360*), *REB1* (*TDEL0D03690*), *SUM1* (*TDEL0E02930*), and *UME6* (*TDEL0F05400*). Ultimately, the tagged alleles were excised from the plasmids by restriction digestion and then integrated into *T. delbrueckii*. The correct integration was confirmed by PCR using primers flanking the sites of recombination. To confirm an intact reading frame, the tagged genes were sequenced, and to confirm protein expression, immunoblots were used.

To tag Abf1, Reb1, and Ume6, the V5 epitope tag and hygromycin resistance gene were amplified from plasmid pFA6a-6xGLY-V5-HphMX4 (FUNAKOSHI AND HOCHSTRASSER 2009) and stitched into the 3' end of each gene (Table S1). To tag Sum1, the V5 tag was generated by site-directed mutagenesis at the 5' end of *SUM1*, and the nourseothricin resistance gene was amplified from pLR1186 (gift from Jasper Rine) and stitched downstream of *SUM1* (Table S1). To tag Kos3, the V5 epitope tag and hygromycin resistance gene were amplified from plasmid pFA6a-6xGLY-V5-HphMX4 and stitched at the 3' end of the gene. Then, a codon-optimized nourseothricin resistance gene was incorporated in place of the HphMX gene using homologous recombination in the *S. cerevisiae* strain JRY4013.

To tag Orc1, a 3xMyc tag was inserted after T248 within the linker between the BAH and $AAA^+$ domains of Orc1, and the hygromycin resistance gene was added downstream of *ORC1* (Table S1). The 3xMyc epitope tag was amplified from plasmid pWZV87 (KNOP *et al.* 1999), and the hygromycin resistance gene was amplified from pFA6a-6xGLY-V5-HphMX4 (FUNAKOSHI AND HOCHSTRASSER

2009). The *orc1-P179A* allele was created by site-directed mutagenesis of the tagged *ORC1* plasmid (Table S1). The *orc1-bah*Δ allele deleted A2 to V213 of *ORC1* and was assembled from three PCR products derived from the tagged *ORC1* plasmid and joined using NEBuilder HiFi DNA assembly master mix (NEB E2621S) (Table S1). All three tagged *ORC1* alleles were integrated into strain LRY3210 with a *KOS3-V5* allele.

To delete *HMR2* or the ORC binding site near *HMR1*, deletion cassettes were created using NEBuilder HiFi DNA assembly master mix (Table S2). The KanMX selectable marker with flanking loxP recombination sites was amplified from pCUG6 (PRIBYLOVA *et al.* 2007) and joined to PCR products corresponding to the sequences flanking the region to be deleted. This assembled deletion cassette was then used to transform yeast. Yeast containing the correct deletion were subsequently transformed with a plasmid expressing Cre recombinase (pZCRE) (PRIBYLOVA *et al.* 2007) to excise the KanMX gene, leaving a single LoxP sequence at the deletion site.

To delete the Rap1 binding sites at *HMR1*, a DNA was synthesized corresponding to the silencer region of *HMR1*, but with the Rap1 binding sites replaced with SalI and SphI restriction sites and a sgRNA cleavage site modified (pLR1358). Strain LRY3208 was transformed with this DNA along with a CRISPR plasmid targeting the *HMR1* silencer (pLR1348, Table S1).

**Yeast transformation**

*T. delbrueckii* cells were transformed using electroporation (HICKMAN AND RUSCHE 2009). Briefly, cells were harvested at an optical density $OD_{600}$ ~1, resuspended in conditioning medium (YPD, 25 mM DTT, 20 mM HEPES, pH 8.0) and incubated for 30 minutes at 30°, and then washed and resuspended in electroporation buffer (270 mM sucrose, 1 mM LiOAc, 10 mM tris, pH 7.5) at 100 OD/ml. 50 µl cells were combined with DNA and electroporated at 300 Ω, 25 µF and 1000 V. Cells recovered 15 minutes in chilled YPD and then four hours at 30° before plating on selective medium (YPD containing 200 µg/mL hygromycin, 300 µg/mL geneticin or 100 µg/mL nourseothricin). For strain construction, 100 ng of linear DNA and 10 μg of sheared salmon sperm DNA (AM9680, Invitrogen) were used. For the *URA3* reporter assay, 50 ng of plasmid was used.

For *S. cerevisiae*, cells were transformed using the polyethylene glycol lithium acetate method (SCHIESTL AND GIETZ 1989). Briefly, cells were harvested at an $OD_{600}$ ~1, washed twice with TEL (100 mM LiOAc, 1 mM EDTA, 10 mM Tris, pH 7.5), and resuspended in TEL at 100 OD/ml. 100 μl cells were combined with DNA and incubated 30 minutes at 30°. Then, 750 μl 40% PEG-TEL was added to each transformation reaction, followed by incubation for 30 minutes at 30° and 10 minutes at 42°. For plasmid construction using homologous recombination, 150 ng of cut plasmid, 300 ng of PCR product, and 10 μg sheared salmon sperm DNA was used.

**Immunoblotting**

Immunoblots were performed as previously described (HICKMAN AND RUSCHE 2010). Cells were grown to an $OD_{600}$ of ~1.0 and fixed with a final concentration of 10% TCA for 20 minutes. For lysis, 45 OD equivalents of fixed cells were vortexed 5 min in the presence of silica beads (0.5 mm dia. #11079105z, BioSpec Products) in 40 μl lysis buffer containing protease inhibitors. Proteins were denatured by the addition of 1/3 volume 3X SDS sample buffer and incubation at 95° for 5 min. Finally, samples were clarified by centrifugation. Aliquots (10 μl) of protein extract were resolved on a 7.5% acrylamide gel, transferred to membrane (Amersham 45004008), and probed to detect Myc-tagged proteins (anti-Myc, Millipore 06-549), V5-tagged proteins (anti-V5, Millipore AB3792), or Pgk1 (ab113687, Abcam) as a loading control.

### SUPPLEMENTAL TABLES

#### Table S1: Plasmids created for yeast strain construction.

| **Cloning procedure** | **(Oligonucleotide number) Sequence [restriction enzyme used]** |
|---|---|
| **pLR1291 (*KOS3*-V5-NatMX)** | |
| *KOS3 (TDEL0E00350)* from JRY10156 ligated to pRS414 (=pLR1290) | (oLR4342) GCGCGccgcggGCACCCAACCTCAGTCTC [SacII] |
| | (oLR4343) GCGCGactagtCTCATCCGCTAGACCTCG [SpeI] |
| V5-HphMX from pFA6a-6xGLY-V5-HphMX4 stitched into pLR1290 (=pLR1376) | (oLR4390) ctaccgcttctttcgacaagcGGGGGAGGCGGGGG |
| | (oLR4391) gtatcagcttcaaatgtccaaaagtggcCTCGTTTTCGACACTGGATGG |
| HphMX in pLR1376 replaced with NatMX (codon optimized gene synthesis) by homologous recombination (=pLR1291) | GCTAGGATACAGTTCTCACATCACATCCGAACATAAACAAatggggactaccctag acgacactgcataccgttaccgtacgtccgtcccaggggatgcagaagccatagaggccttagacggatcgttcaccacg gatactgtgttccgtgtcacagccacgggcgatggtttcactcttagagaagtacccgttgatccgccgcttactaaagtattt ccagatgatgaatcagatgatgagtcagacgacggagaggatggcgacccggacagtaggacgttcgttgcttatggag acgacggcgacttggccgggtttgtagtcgtgtcttactcagggtggaaccgtagattaaccgtagaagacatagaagtcg ctcccgagcatagagggcacggagtcgggcgtgctttgatgggcttagctaccgagtttgctagggagcgtggcgccgg gcacctatggcttgaggtaactaacgtaaatgcaccagctattcacgcttatagaaggatggggtttacattgtgtggacttg ataccgctctatacgatgggactgcttcagatggggaacaagctctttatatgagtatgccgtgtcccTCAGTACTG ACAATAAAAAGATTCTTGTTTTCAAGAACTT |
| **pLR1289 (*ORC1*-Myc-HphMX)** | |
| *ORC1 (TDEL0D00910)* from JRY10156 ligated to pRS414 (=pLR1187) | (oLR4251) GCGCGccgcggCCAGGACAAGGGCAGAGATC [SacII] |
| | (oLR4252) CAGCAGGAACTGCCGAAG [endogenous SpeI] |
| HphMX from pFA6a-6xGLY-V5-HphMX4 stitched into pLR1187 (=pLR1339) | (oLR4415) gaacaggatgaaagtttgaaaggtctatgaGTAGGTCAGGTTGCTTTCTCAG |
| | (oLR4416) gaagaagttcaccatctacgcgCTCGTTTTCGACACTGGATGG |
| 3xMyc from pWZV87 stitched into pLR1339 (=pLR1289) | (oLR4874) cataaaattggagcccgacacgGCTGCTAGTGGTGAACAAAAG |
| | (oLR4875) gaatcttcgctggcactatcGGATCCGTTCAAGTCTTCTTC |
| **pLR1293 (*orc1-P179A*-Myc-HphMX)** | |
| *P179A* mutation [CCT>GCT] made by site directed mutagenesis in pLR1289 (=pLR1292) | (oLR4713) CAAGGATTTCTTAGTAAGATATATCTGTGAGgctACCGGTGAGAATTTTGCTC |
| | (oLR4714) GAGCAAAATTCTCACCGGTagcCTCACAGATATATCTTACTAAGAAATCCTTG |
| **pLR1332 (*orc1-bahΔ*-Myc-HphMX)** | |
| PCR products made using oLR4997/4998, oLR4995/5010 and oLR5009/4996 with pLR1289 template assembled using NEBuilder HiFi Mix (=pLR1332) | (oLR4997) ccgatcgaagtctgagtgtttgaaacatgCCGAAGTCTTTGAACTCGC |
| | (oLR4998) GATCTGCCGGTAGAGGTG |
| | (oLR4995) GTAGGTCAGGTTGCTTTCTCAG |
| | (oLR5010) CTCGAGGTCGACGGTATC |
| | (oLR5009) GATACCGTCGACCTCGAG |
| | (oLR4996) CATGTTTCAAACACTCAGACTTCG |

| | |
|---|---|
| **pLR1302 (*ABF1*-V5-HphMX)** | |
| *ABF1 (TDEL0A02360)* from JRY10156 ligated to pRS414 (=pLR1299) | oLR5267 GCGCGactagtCTCTGGCATTGACATTCCC [SpeI] |
| | oLR5327 GCGCGccgcggGTCGATGCCTTTCACAGCAC [SacII] |
| V5-HphMX from pFA6a-6xGLY-V5-HphMX4 stitched into pLR1299 (=pLR1302) | oLR5341 gaacattcagccagaattgagaggtcaaGGGGGAGGCGGGGG |
| | oLR5342 gacattacttggagatttttcaatatcaaattgttcgcCTCGTTTTCGACACTGGATGG |
| **pLR1303 (*REB1*-V5-HphMX)** | |
| *REB1 (TDEL0D03690)* from JRY10156 ligated to pRS414 (=pLR1300) | oLR5270 GCGCGccgcggCTGTAACCCAACCTGACAAGAC [SacII] |
| | oLR5271 GCGCGactagtCAAGACCCCAAAGACCGC [SpeI] |
| V5-HphMX from pFA6a-6xGLY-V5-HphMX4 stitched into pLR1300 (=pLR1303) | oLR5297 caattccactagattccaaaggtaatGGGGGAGGCGGGGG |
| | oLR5298 caaaatttgccctgttgatcatacCTCGTTTTCGACACTGGATGG |
| **pLR1304 (*UME6*-V5-HphMX)** | |
| *UME6* (*TDEL0F05400*) from JRY10156 ligated to pRS414 (=pLR1301) | oLR5268 GCGCGcccgggCCACATCAACCCACGAAGTAC [XmaI] |
| | oLR5269 GCGCGggtaccCGAGTCAATAGCTCCGTCG [KpnI] |
| V5-HphMX from pFA6a-6xGLY-V5-HphMX4 stitched into pLR1301 (=pLR1304) | oLR5295 gaaaccaactcgatccagttccGGGGGAGGCGGGGG |
| | oLR5296 cctagatggattcttcctcatcgtgCTCGTTTTCGACACTGGATGG |
| **pLR1330 (V5-*SUM1*-NatMX)** | |
| *SUM1 (TDEL0E02930)* from JRY10156 ligated to pRS414 (=pLR1328) | (oLR4458) GCAGTTCCATGAGACAAGACAC [endogenous EcoRI] |
| | (oLR4459) GCGCGccgcggGGCCCTTAGATGGGTCATC [SacII] |
| NatMX from pLR1186 stitched into pLR1328 (=pLR1329) | (oLR4492) gtcatcccacccacgcaatagGTAGGTCAGGTTGCTTTCTCAG |
| | (oLR4495) ccattgcttcatcgttccagtcgCAGCAGTATAGCGACCAGC |
| V5 tag added by site directed mutagenesis to pLR1329 (=pLR1330) | (oLR4510) GAGTATCAGGGTGATTTGTAAGATGggaaagcctatccctaaccctctcctcggtctcgattctacgATCCAGGGTATAAGGGCTAGTTC |
| | (oLR4511) GAACTAGCCCTTATACCCTGGATcgtagaatcgagaccgaggagagggttagggataggctttccCATCTTACAAATCACCCTGATACTC |
| **pLR1348 (Cas9 + sgRNA targeting *HMR1* near Rap1 binding sites)** | |
| HiFi assembly of oLR5422 and NotI-cut pXIPHOS-panARS | (oLR5422) CGGGTGGCGAATGGGACTTTcgttcagataagttttggggGTTTTAGAGCTAGAAATAGC |
| **pLR1358 (*HMR1* silencer with Rap1 binding sites deleted (Rap1$_{bs}\Delta$))** | |
| Gene synthesis product with Rap1 binding sites | GAGAGTCGTATGTGGAGCAAAACAGCTCAAAGTCACCTGACAAAGTTCATGATCGCTAGCGATGTATATAGTCGCCATACACGAGTATTATCGTTTACTTG |

| | |
|---|---|
| near *HMR1* replaced with SalI and SphI restriction sites and modified sgRNA-directed cleavage site | TCTAAACACTCGTAGGTCATGATATTAGAAGTAAATATTCGTATTGAAGCACAGTACACTGCTGTAAATACTCGTCATGTATGAGCAAAGAAGGAGCGGGTAACAAACTACTACAACGTACTGGGCaTTCAGATAAaTTTTGaGGtttAATAACAATAATTTCAAGGCGAGGTGATTATCATGCTTCAGAATgtcgacATGGTAATAAATGTAGATTCTAGAACAGTAGTATTTGAGAGACCGGTCAGCCTTGAATACCAGCAGCCTTTATTTAGTGCTTACTCCCTATTATTACATCTGGGTGAATATGGGAACAATCCAAGTACTGACTACTATATGTTCATAAACACATATATAATGAAGCAAAAGTAACGTGAGTAAACTTATTGTATATAAGTCTGTCAAATATTAGTTTTACATCAAAATTTGTTGGTTTCGTGGTCTAGTTGGTTATGGCATCTGCTTAACACGCAGAACGTCCCCAGTTCGATCCTGGGCGAAATCAATTTTTTTCGAAAgcatgcTAATACAGAAATATCCAAGTACACCACAGCGGCTTGAGACAGAACGAACTCGTCCATGTCGCGACAATAAATAAAGCCTCTGACAAAACTAGGTTTGAGAGCAACATCACGTTGATCGAAAAGTTGTTAGCCACTGCACAATTGAATATATATTACCAGATGTTAAAACAGTCTGACCCGCTGGCCGTTTTTTTCAGCTGGCATCATCCAGTTTG |

**Table S2: Oligonucleotides used for strain construction.** Deletion cassettes were created by combining three PCR products using NEBuilder HiFi DNA assembly master mix.

| Allele | Set | Oligo | Sequence |
|---|---|---|---|
| *hmr2Δ::LoxP* | KanMX & LoxP | oLR5047 | gaaagagagctccaagccgttcAAGCTTCGTACGCTGCAGGTC |
| | | oLR5048 | gaaaaagggcactgcaaaatgtcaagcggGACTCACTATAGGGAGACCG |
| | *HMR2* silencer | oLR5051 | CGTGCCAAATGATACCGCC |
| | | oLR5052 | CTTGAACGGCTTGGAGCTCTC |
| | flanking *HMR2* | oLR5049 | CCGCTTGACATTTTGCAGTGC |
| | | oLR5050 | GGAGCCCAGACCTGTAAC |
| *arsVII22Δ::LoxP* | KanMX & LoxP | oLR5272 | gtctcaaagccacaattgacaacctacttGCTTCGTACGCTGCAGGTC |
| | | oLR5340 | catgtttgaagcttttgtcacacgcacgaggGACTCACTATAGGGAGACCG |
| | Up of 0G00140 | oLR5274 | CCGTAAGGATTAAGTGGCAC |
| | | oLR5275 | CATCGCAATTCAAGTAGGTTGTC |
| | Down of 0G00150 | oLR5339 | GGTTTCCTCGTGCGTGTGAC |
| | | oLR5277 | CTCATGCGGCTGAGATCG |

**Table S3: Plasmids for *URA3* silencing assay.** All plasmids except for empty vector include *TdHMR2* with **a**1 open reading frame replaced with *K. lactis URA3*.

| Cloning procedure | Oligonucleotide and Sequence |
|---|---|
| **pRS41K-Td*CEN3* (Empty Vector)** | |
| *T. delbrueckii* shuttle vector with KanMX | From (ELLAHI AND RINE 2016) |
| **pLR1377 (*HMR2* silencer)** | |
| Replace HphMX in pRS41H-Td*CEN3*-***a**1Δ::URA3* with KanMX from p3FLAG-KanMX using NEBuilder HiFi mix | oLR5533 gattgtactgagagtgcagcgaCATGGAGGCCCAGAATACCC |
| | oLR5537 cggtatttcacaccgccgtcccGAGCTCGTTTTCGACACTGG |
| | pRS41H-Td*CEN3*-***a**1Δ::URA3* cut with MluI, excising HphMX gene |
| **pLR1382 (*HMR2* silencer lacking ORC binding site (*arsV75Δ*))** | |
| TdARS-V75 replaced with SphI site by mutagenesis of pLR1377 | oLR5525 CCCCTAATTTATTGAATTGAACCACgcatgcCCAAGATTTATTTAAAACCACACCG |
| | oLR5526 CGGTGTGGTTTTAAATAAATCTTGGgcatgcGTGGTTCAATTCAATAAATTAGGGG |
| **pLR1384 (*HMR1* silencer)** | |
| *HMR1* silencer from JRY10156 replaces *HMR2* silencer in pLR1377 by stitching | oLR5574 gcgcaattaaccctcactaaaggGAACGGACCCATTGAGCAG |
| | oLR5575 ggtcaactggatgatggtcGGCGAAGAGATATTCAGGC |
| **pLR1399 (*HMR2* silencer lacking ORC and Abf1 binding sites)** | |
| ORC & Abf1 binding sites replaced with SphI by mutagenesis of pLR1377 | oLR5645 GCATCCAGTGAGATTCGCgcatgcCATGAGTGACTTACGGCAC |
| | oLR5646 GTGCCGTAAGTCACTCATGgcatgcGCGAATCTCACTGGATGC |
| **pLR1400 (*HMR2* silencer lacking ORC binding site (*arsV75Δ*); plasmid origin deleted (*ars209Δ*))** | |
| *ScARS209* replaced with SphI & NheI sites by mutagenesis of pLR1382 | oLR4990 GGTTTCTTAGGACGGATCGgcatgcaatcgggctagcGATCCCCCTAGAGTCTAAGC |
| | oLR4991 GCTTAGACTCTAGGGGGATCgctagcccgattgcatgcCGATCCGTCCTAAGAAACC |
| **pLR1401 (*HMR2* silencer; plasmid origin deleted (*ars209Δ*))** | |
| *ScARS209* replaced with SphI & NheI sites by mutagenesis of pLR1377 | oLR4990 GGTTTCTTAGGACGGATCGgcatgcaatcgggctagcGATCCCCCTAGAGTCTAAGC |
| | oLR4991 GCTTAGACTCTAGGGGGATCgctagcccgattgcatgcCGATCCGTCCTAAGAAACC |

**Table S4: Genes used to construct species tree**

| *S. cerevisiae* gene | Description |
|---|---|
| *YBR202W;* NP_009761 | Mcm7, ATP-dependent DNA helicase component of the MCM mini-chromosome maintenance complex involved in initiation and regulation of DNA replication |
| *YPL086C;* NP_015239 | Elp3, Subunit of Elongator complex required for modification of wobble nucleosides in tRNA |
| *YHR186C;* NP_012056 | Kog1, Subunit of the TORC1 complex; binds to ubiquitin |
| *YDL215C;* NP_010066 | Gdh2, Mitochondrial glutamate dehydrogenase involved in nitrogen metabolism |
| *YMR108W;* NP_013826 | Ilv2, Acetolactate synthase: protein involved in the biosynthesis of branched chain amino acids (isoleucine, valine and leucine) |
| *YKL210W;* NP_012712 | Uba1, Ubiquitin activating enzyme (E1) |
| *YNL102W;* NP_014247 | Pol1, Catalytic subunit of the DNA polymerase I alpha-primase complex |
| *YJL008C;* NP_012526 | Cct8, Subunit of the cytosolic chaperonin Cct ring complex |
| *YDR190C;* NP_010476 | Rvb1, ATP-dependent DNA helicase |
| *YCR092C;* NP_010016 | Msh3, Mismatch repair protein |
